# Substituent-based Modulation of Self-Assembly and Immunogenicity of Amphipathic Peptides

**DOI:** 10.1101/2025.07.08.663637

**Authors:** Anirban Das, Ushasi Pramanik, Elise M. Brown, Chih-Yun Liu, Huan Gong, Jonathan Fascetti, Mark Gibson, Samuel Stealey, Silviya P. Zustiak, Cory Berkland, Meredith E. Jackrel, Mark A. White, Jai S. Rudra

## Abstract

Peptide-based biomaterials assembled through monomer-by-monomer self-assembly provide versatile platforms for biomedical applications due to their adjustable physicochemical properties, biocompatibility, and dynamic nature. The self-assembly process largely depends on primary sequence features, such as hydrophobicity, length, and charge, which influence the formation of various nanostructures, including fibrils and hydrogels. Amphipathic peptides, characterized by alternating polar and hydrophobic residues, are especially effective in forming supramolecular nanofibers stabilized by π–π interactions and hydrogen bonds. Chemical modifications, particularly on aromatic side chains, have proven to be a promising approach for controlling assembly morphology, stability, and biological activity. In organic chemistry, the use of chemical substituents, such as halogens, alkyl groups, or electron-donating and electron-withdrawing groups, has been widely employed to alter reactivity, stability, and molecular interactions for diverse applications, including catalysts, pharmaceuticals, and materials science. However, the influence of these substituents on peptide packing and *in vivo* immunogenicity remains relatively unexplored. In this study, we systematically examine how changes in the position and nature of substituents on benzyl groups attached to short amphipathic peptides affect self-assembly, fibril morphology, and immune responses. By introducing different electron-donating and withdrawing groups at the para-position of benzyl rings and modifying the chain length connecting the backbone to the aromatic moiety, we observe notable effects on fibril formation, molecular packing, and immunogenicity both *in vitro* and *in vivo*. Our results show that subtle chemical modifications are effective tools for designing tailored peptide nanomaterials with promising potential in vaccine delivery, tissue engineering, and regenerative medicine.

## INTRODUCTION

Peptide-based biomaterials developed through monomer-by-monomer self-assembly offer significant advantages due to their unique physicochemical properties, biocompatibility, and dynamic characteristics.^1^ Peptides serve as ideal building blocks due to the chemical diversity of amino acids, allowing for precise control over self-assembly triggers such as pH, temperature, and ionic strength.^2–4^ By tailoring primary sequence features, including hydrophobicity, length, chirality, and charge, various superstructures, such as micelles,^5^ nanofibers,^6–9^ nanotubes,^10–12^ and vesicles,^13^ can be engineered to possess specific properties. Amphipathic designs with alternating polar/charged and hydrophobic residues readily self-assemble into high-aspect-ratio cross-β fibril nanofibers in aqueous buffers.^14^ These fibers entangle at sufficiently high concentrations to form self-supporting hydrogels with numerous biomedical applications. In particular, (FKFE)_n_ repeat sequences featuring phenylalanine (F) as the nonpolar component are well-characterized, with abundant data relating to sequence length and variation patterns of hydrophobic and charged residues on self-assembly.^15,16^ The minimal length, strong assembly propensity, and gelation properties of KFE8 (double repeat of FKFE) have made it the most thoroughly studied member of this peptide class.^17–19^

The stability of supramolecular peptide assemblies relies on a delicate balance of non-covalent interactions, primarily hydrogen bonding, π-π stacking, electrostatic, van der Waals forces, steric repulsions, and hydrophobic interactions.^20,21^ Studies show that removing a single terminal residue from KFE8 significantly alters and, under certain conditions, completely prevents self-assembly. In recent years, employing substituent groups (such as halogens, alkyls, and alcohols) to disrupt this equilibrium and promote assemblies with diverse scales and morphologies has emerged as a significant area of research in materials science and engineering.^4,22^ Substituent effects are also of great interest to chemists, finding applications in supramolecular chemistry to modulate the structures of metal coordination cages, polymers, and metal-organic frameworks (MOFs).^23,24^ Notably, substitutions involving activating electron-donating groups (EDGs) or deactivating electron-withdrawing groups (EWGs) on benzyl rings have been extensively studied in organic chemistry. The subtle changes in electron cloud distribution due to substituent effects significantly impact the reactivity, stability, and biological activity of the molecule, thereby influencing its structure and interactions with other molecules. Furthermore, the position of the substituent on the ring (ortho, meta, or para) can affect the packing structure and the molecules’ orientation.

Given the essential role of hydrophobic residues in the self-assembly of amphipathic peptides, Nilsson and colleagues leveraged the abundance of benzyl groups in (FKFE)_n_ peptides to create a valuable framework for understanding how substituent groups such as CH_3_, Cl, Br, F, OH, NO_2_, and CN influence π–π interactions in self-assembly or co-assembly processes.^4^ However, prior studies relied on minimal systems consisting of a single F residue or diphenylalanine (FF) peptides linked to a fluorenyl methoxycarbonyl (Fmoc) protecting group.^25,26^ The reported crystal structure of Fmoc-FF by Adams and coworkers shows that π-stacking interactions between Fmoc groups and side chain phenyl groups, along with the hydrogen bonding interactions among carbamate and amide groups, are key drivers of self-assembly.^27^ Other studies using natural amyloid sequences (KLVFF) or designed peptides (FQFQFK) focused solely on halogenation to modulate assembly kinetics, morphology, and gel stiffness. A significant knowledge gap exists in understanding how substituents on the benzyl group influence the assembly and molecular packing of F-rich peptides that lack Fmoc protecting groups. Moreover, no studies have reported the *in vivo* effects of substituent modifications on the (FKFE)_n_ class of peptides, making it crucial to address this gap, as insights from these studies could lead to the development of higher-order and multi-component hydrogels for biomedical applications.

**Scheme 1.**
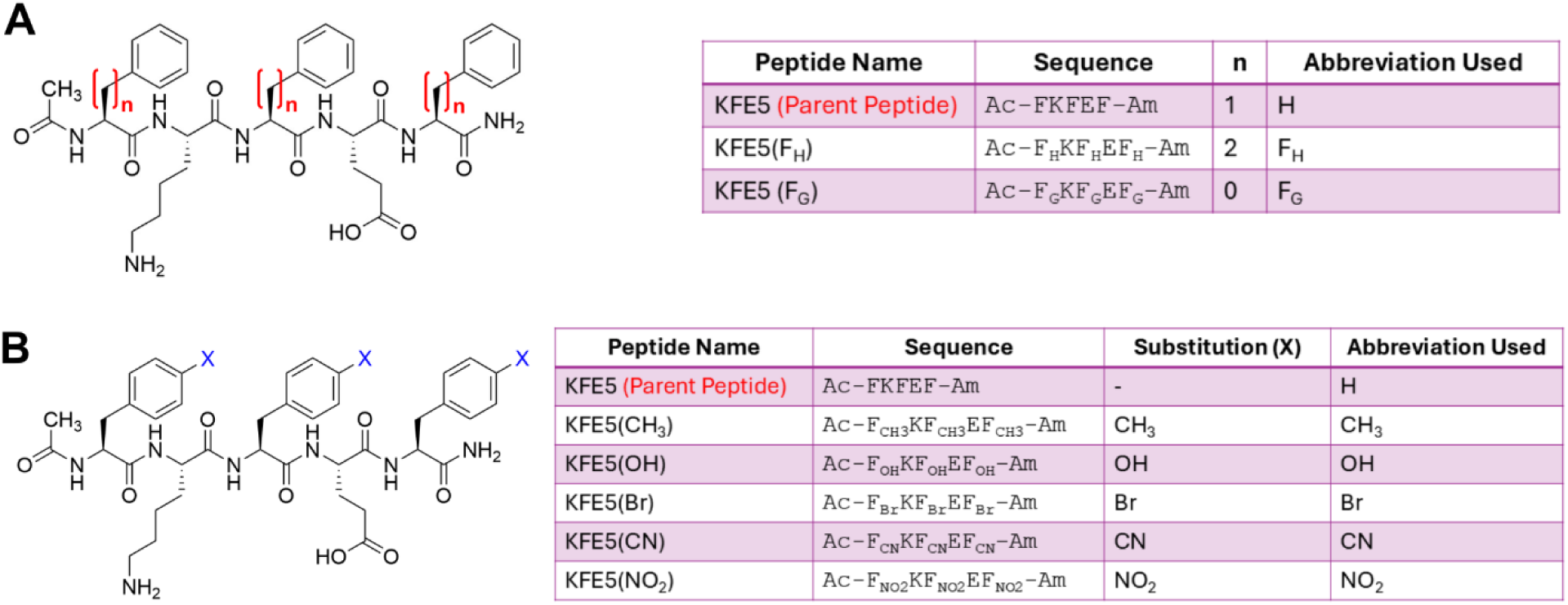
Sequences of KFE5 (A) positional variants and (B) chemical variants used in this study.

In this study, we utilized KFE5 (Ac-FKFEF-Am) (**Scheme 1**), a truncated variant of KFE8 (Ac-FKFEFKFE-Am), to investigate the effects of substituents on the self-assembly and immunogenicity of supramolecular peptide nanofibers. KFE5 was identified as the minimal sequence necessary for self-assembly, as no fibril formation was observed with KFE4 (FKFE). We also varied the distance of the benzyl group from the backbone by replacing F with either the more rigid phenylglycine (F_G_) or the more flexible homophenylalanine (F_H_). This resulted in the benzyl group being placed at zero carbon distance (n=0, F_G_), one carbon distance (n=1, F), or two carbon distance (n=2, F_H_) from the peptide backbone (**Scheme 1A**). We introduced EWGs (Br, CN, NO_2_) or EDGs (CH_3_, OH) at the para position (**Scheme 1B**) to assess their impact on self-assembly. Thorough biophysical and biochemical characterization utilizing various spectroscopic techniques revealed a fundamental β-sheet-rich fibrillar nature with diverse molecular packing modes, morphologies, and bulk rheological properties. Cross-seeding assays with FRET biosensor cells demonstrated that the constructs are non-toxic and non-amyloidogenic. Additionally, treating primary bone marrow-derived dendritic cells (DCs) with modified nanofibers resulted in differing innate immune activity. RNA-seq analyses in primary DCs indicated upregulated stress response pathways, and all constructs were compatible with modification using the model antigenic peptide OVA_323-339_ (ISQAVHAAHAEINEAGR). In mice, the antibody and cellular immune responses developed in a substituent-dependent manner. Our findings suggest that substituents offer a wider range of design blocks for constructing synthetic peptide assemblies, serving as a straightforward yet potent tool for creating custom biomaterials in engineering and medicine.

## MATERIALS AND METHODS

### Peptide Synthesis

Standard Fmoc-SPPS chemistry was used to synthesize the peptides on Rink Amide resin. Oxyma [ethyl 2-cyano-2-(hydroxyimino)acetate] and *N,N′*-Diisopropylcarbodiimide (DIC) were used as coupling agents on a Liberty Blue microwave-assisted synthesizer. Peptide cleavage was carried out in a cocktail of trifluoroacetic acid (TFA), triisopropylsilane (TIS), and H_2_O (95:2.5:2.5) and diethyl ether was used for extraction. The resulting pellet was frozen and lyophilized in acetonitrile/water (ACN/H_2_O) mixture (50:50). Peptides were purified (∼90%) using High Performance Liquid Chromatography (HPLC) on a Dionex Ultimate 3000 HPLC equipped with a diode-array detector. ACN/H_2_O gradient was applied to an Agilent Poroshell RP-C18 column (4.6 mm×150 mm), operating at a 1 mL/min flow rate. MALDI-TOF mass analysis (Shimadzu MALDI-8030) was used to confirm peptide identity using α-cyno-4-hydroxycinnamic acid matrix (Bruker Daltonics, MA) **(Figures. S1-S8)**. OVA_323-339_ conjugated peptides were purchased from GenScript or P3 Bio-Systems and used without further modification.

### Peptide Fibrillization

Peptide self-assembly was achieved by dissolving the purified peptide in Ultrapure Water (UPW) to prepare 1 mM stocks for further dilution and use. OVA-conjugated peptides were first dissolved in UPW, then diluted in 10× PBS to achieve a final concentration of 1 mM in 1× PBS for immunizations.

### Peptide Modeling and Molecular Dynamics

The FKFEF peptide, excluding the peptide modifications, was used in AlphaFold3 with 30 copies. The most complete and continuous model was extended to a full 16-strand sheet of the predicted 2-start fiber with edge contacts between the two parallel fibers using COOT and PyMol. Each fiber was assembled similarly to that in KFE8,^32^ with the phenylalanine residues forming a hydrophobic core sandwiched between the peptide β-sheet forming backbone. This protofiber model, with the N-acetyl and amidated C-terminus modifications added, was allowed to equilibrate in explicit TIP3 solvent with 150 mM NaCl, using NAMD3 with initial annealing, with anti-parallel β-sheet hydrogen binding restraints applied using the extra bonds option, followed by a 500 ns full free MD simulation. The resulting protofilament model was then extended to a 48-strand sheet, 2-start fiber, dual parallel fiber bundle of 192 KFE5 molecules. This extended model was minimized and annealed with model anti-parallel β-sheet hydrogen bonding restraints as for the protofilament model. The 192-chain model was then equilibrated for 50 ns at 300 K.

### Thioflavin T (ThT) Assay

For ThT assays, 50 µM fibrillated peptide solutions were mixed with 20 µM ThT solution, and fluorescence was monitored at 485 nm upon excitation at 440 nm in a 96-well plate using a Synergy HT plate reader (Biotek, USA).

### Transmission Electron Microscopy (TEM)

Peptide solutions (10 μM) were applied directly to 200-mesh, carbon-coated copper grids for 2 min and stained with 1% uranyl formate for 1 min. Excess stain and the peptide solution were blotted using filter paper, and grids were washed thrice with UPW. Darkfield images were acquired on a JEOL JEM-2100F Field-Emission STEM microscope at an accelerating voltage of 120 kV.

### Width Measurements of Fibrils

FIJI software was utilized to measure the widths of the fibers, and the measurement scale was established using the scaling factor displayed in each image. Individual fibers were selected for measurement, avoiding aggregates. The width of each fiber was measured using the line tool by drawing a line across the fiber’s width at an angle approximately perpendicular (90°) to its orientation to ensure accuracy. The start and end points of the line were determined based on the contrast between the fiber and the surrounding background. Approximately 30 measurements were randomly taken for each sample.

### Fourier Transform Infrared (FT-IR) Spectroscopy

FT-IR measurements for secondary structural analysis were conducted using 1 mM peptide solutions (1×PBS, pH 7.4) with a Bruker Alpha II FT-IR instrument equipped with a Smart Performer single-reflection ATR accessory and an Au crystal sample stage. The background and buffer spectrum were collected and subtracted from the sample using OPUS software. An average of 24 scans for each peptide were employed for each sample measurement. Data were analyzed using GRAMS/AI software (Thermo Scientific, USA). Second derivative spectra were calculated from the absorbance spectra in the Amide I region using a Savitzky-Golay filter, third order, with a nine-point window. The second derivative spectra analyzed between 1610 cm^-1^ and 1710 cm^-1^ were fit with six or seven Gaussian curves, informed by the Akaike information criterion, and the peak positions were compared to literature reports.^33–35^

### Circular Dichroism (CD) Spectroscopy

CD spectra were recorded using 100 μM peptide solutions on a Jasco J-815 CD spectrometer at RT (25°C). The spectra (average of three scans for each sample) were collected using the following parameters: wavelength 190–260 nm, bandwidth of 1 nm, data pitch of 0.5 nm step. The solvent background was subtracted from each spectrum.

### Mechanical Properties of Peptides

An ARES 2000ex rotational rheometer (TA Instruments, New Castle, DE) was used to evaluate the viscosity and viscoelastic properties of the peptides. Peptides were freshly solubilized in deionized water at 5 mM and allowed to incubate for 10 min to allow for complete solubilization. Next, 170 µL of each peptide solution was pipetted directly onto the rheometer stage. A 20 mm parallel plate geometry was lowered to a gap of 200 µm. Storage modulus (G′) and loss modulus (G″) were measured as a function of strain amplitude of 0.05-50% at an angular frequency of 10 rad/s to determine the linear viscoelastic region. Next, G′ and G″ were measured as a function of angular frequency (1-10 rad/s) at a strain of 1%. Lastly, the viscosity of each peptide solution was measured as a function of shear rate (0.1-100 s^-1^) to determine if samples exhibited non-Newtonian behavior. Separately, a Hagen-Poiseuille viscometer (microVISC, Rheosense, San Ramon, CA) was used at a shear rate of 1000 s^-1^ to validate measured viscosity trends.

### X-ray Scattering Studies

Small-angle X-ray scattering (SAXS) data of the KFE5 X-phenyl substituted peptide solutions were collected using a Rigaku (Woodlands, TX) BioSAXS-1000 camera on an FRE^++^ X-ray generator with an ASC-96 Automated Sample Changer held at 10°C. A matching buffer was collected for each sample. The detector was calibrated using a silver behenate powder sample, following the manufacturer’s recommended procedure. The SAXS samples (KFE5 and its variants) were prepared by dissolving 2-3 mg of lyophilized peptide in 400 µL of deionized water to produce a final concentration of 4 mg/mL. The samples were then vortexed, and precipitants were removed by centrifugation at 13.3 rpm for 5 minutes. Although the samples formed gels, these remained fluid under hydrostatic pressure, permitting pipetting by the ASC-96 liquid handling system. The cyano variant of KFE5 was an exception. It remained a gel at 4 mg/mL and needed to be diluted to 2 mg/mL to be pipetted or flowed in the ASC-96 liquid sample handling system. Processing was performed in SAXSLab (Rigaku) and SAXNS-ES (https://xray.utmb.edu/SAXNS). Analyses were performed in Primus/GNOM,^36^ BIFT,^37,38^ and gnuplot (http://www.gnuplot.info).

WAXS samples were prepared by dissolving 3-5 mg of lyophilized peptide in 500 µL of methanol. Each sample was pipetted into MiTeGen capillary in 30 µL aliquots, left to dry, and additional aliquots added until the entire sample was transferred into the capillary. Powder diffraction data were collected on a Rigaku R-AXIS-IV^++^, with a Rigaku FRE^++^ Cu X-ray source using four 30-minute frames. The diffraction images were processed using Fit2D and calibrated with a sucrose sample. The curves from multiple frames were averaged in Primus. In addition to the strong powder diffraction, the data contained broad background scattering from the capillary and the amorphous gel. This background was removed using d1Dplot and the resulting powder diffraction peaks fit. X-ray diffraction peaks were each fit to a simple Gaussian with a common theta-dependent peak width σ using gnuplot.

### Bone Marrow-Derived Dendritic Cells (DCs) Culture

The femurs and tibiae of mice of C57BL/6 mice were flushed, and single-cell suspensions were prepared, following removal of red blood cells (RBCs) using lysis buffer (Invitrogen, USA). After washing twice using HBSS (Hanks balanced salt solution), cells were differentiated for 7-9 days in complete RPMI1640 media containing heat-inactivated FBS (10%), β-mercaptoethanol (55 µM), Sodium pyruvate (1 mM), HEPES (10 mM), 1× MEM Non-essential Amino acids, 20 ng/mL granulocyte-macrophage colony-stimulating factor (GM-CSF), 5 ng/mL IL-4 and 100 μg/mL Penicillin-Streptomycin (pen-strep) in 150 mm (25 mL/dish) in an incubator at standard conditions (37°C and 5% CO_2_). Cells were harvested using gentle washing between days 7-9 for experiments.

### Antigen Presentation Assays

To assess antigen presentation, DCs (5×10^4^ cells/well) were plated in 96-well plates and treated with OVA-conjugated peptides at various concentrations (0.1, 1, 2.5, 5, 10 μM) for 24h. The cells were then washed with PBS to remove extracellular fibrils. The T-cell hybridoma cell line DOBW (kind gift from Dr. Clifford V. Harding, Case Western Reserve University, Cleveland, OH, USA)^39^ were overlaid (1:5, DC: hybridoma) for 16h. The supernatant was collected, and IL-2 levels were quantified by ELISA (Biotechne, Cat#: DY402-05). Briefly, 96-well high-binding ELISA plates (CORNING Costar 3361) were coated with 100 µL of the capture antibody in 1× PBS (pH 7.4) overnight at 4°C and then blocked with blocking buffer (1% BSA in 1× PBS, pH 7.4) for 1h. After the blocking step, cell culture supernatants or standards were added to the wells in a diluent buffer (0.1% BSA and 1× TBST) and incubated for two hours, followed by the addition of a detection antibody (100 µL/well in the diluent buffer for 2h). Finally, streptavidin-HRP (100 µL/well) in diluent buffer was added for 20 minutes, followed by 100 µL of TMB (Tetramethylbenzidine) substrate. The reaction was stopped using 50 µL of 1 M phosphoric acid, and absorbance was measured at 450 nm. The plate was washed 3× using 300 µL/well of wash buffer between each incubation step. The amount of IL-2 produced was measured from the standard curve.

### Immunizations and Antibody Titers

All experiments were approved by the Institutional Animal Care and Use Committee (IACUC) at WashU. Female C57BL/6 mice (Jackson Labs 4-6 weeks old) were housed under standard conditions. Mice (n = 6/group) were subcutaneously (s.c.) vaccinated (100 µL of 2 mM solution) and boosted three weeks later with the same dose. Control mice received saline. 7-10 days post-boost, blood was collected using cardiac puncture, and serum was extracted by centrifugation. High-binding ELISA plates (Corning #3361) were coated with 20 μg/mL OVA peptide in PBS and incubated overnight at 4°C and subsequently blocked for 1h at RT with 1× ELISA diluent (Invitrogen; 100 μL/well). After blocking, 50-fold dilutions of the sera were prepared in ELISA diluent and added to the wells (100 μL/well for 2h at 25°C) followed by a secondary HRP-conjugated goat anti-mouse IgG (1:5000, 100 μL/well) for 30 min. For isotyping, HRP-conjugated isotypes (IgG1, IgG2c; Southern Biotech) were added (1:4000, 100 μL/well for 30 min at 25°C) after the addition of sera prior to development. Plates were developed using TMB substrate (100 μL/well) for 15 min, and the reaction was quenched with 1 M phosphoric acid (50 μL/well). The plate was washed 3× using 300 µL/well of wash buffer between each incubation step. Absorbance was recorded at 450 nm using a BioTek Synergy microplate reader, and background absorbance from non-antigen-coated wells was subtracted.

### Cellular Immune Responses through Flow Cytometry Analysis

Following euthanasia, spleens were dissociated using syringe plungers through 100 µm filters and washed with HBSS containing 5% FBS. Cell pellets were resuspended in 2 mL RBC lysis buffer for 2 min, washed with HBSS, and resuspended in complete RPMI 1640. Cells were counted and plated in triplicate (10^6^ cells/well in 250 µL), and antigen-specific T cell populations were recalled by incubating with RPMI 1640 media containing 0.2 mg/mL of the cognate antigen OVA. After 96 hours, cells were washed twice with 200 µL PBS, then resuspended in 100 µL PBS containing eFluor506 (eBioscience, 65-0866-14) and Fc-Block (BioLegend, 101302) for 30 minutes. Cells were then washed 2× with FACS wash buffer (FWB, 1× PBS + 10% FBS) and stained with 50 µL FWB with OVA-specific Tetramer (I-A^b^ PE, MBL TS-M710-1), 50 µL of anti-CD3 (BV421, Biolegend 100228), and 50 µL of anti-CD4 (PerCP-Cy5.5, eBioscience, 45-0042-82) for 30 min. Cells were washed 2× with FWB and resuspended in 4% paraformaldehyde for fixation. After 15 minutes, cells were washed three times and resuspended in 200 µL FWB for acquisition. Cells were analyzed on a Beckman Coulter CytoFLEX S, and data were analyzed using FlowJo software. Single-color control beads were used for compensation. For analysis, cells were gated on single cells, live, CD3^+^, CD4^+,^ and tetramer^+^ populations.

### RNA Sequencing

Mature BMDCs (5×10^5^ cells/well) were plated in 24-well plates and treated with 10 μM peptide solutions for 24 h, following which, the QIAGEN RNeasy Mini Kit (Cat#: 74104) was used for the isolation and purification of RNA from the treated DCs. Briefly, 350 μL of Buffer RLT was added to the pelleted cells and vortexed for complete homogenization. It was then followed by an addition of 350 μL of 70% lysate and mixed well by pipetting. The entire volume of the sample was then transferred to a RNeasy Mini spin column placed in a 2 mL collection tube. The tube was centrifuged for 20 s at 8000× g at 4°C. The flow through was discarded and 700 μL of Buffer RW1 was added, following a 20 s 8000 × g centrifugation at 4°C. Similarly, after discarding the flow-through, the spin column was centrifuged under similar condition by the addition of 500 μL of Buffer RPE 2× times. The RNA spin column was placed in a new 1.5 mL collection tube followed by addition of 50 μL of RNase-free water. The solution was centrifuged at above mentioned conditions, and the RNA yield was estimated for each sample using the Nucleic Acid Quantification Protocol in a BioTek Synergy H1 Microplate Reader.

The mRNA library construction, quantification, and sequencing via Illumina platforms were performed by Novogen (Sacramento, CA, USA) with stepwise quality control checks, which includes read counts normalization, model dependent p-value estimation, and FDR value estimation based on multiple hypothesis testing. This is preceded by filtering raw reads, mapping clean reads to a reference genome using HISAT2, and determining FPKM values for all samples. Differentially expressed genes were evaluated based on their log_2_ fold change and adjusted p-values. Those with |log_2_(fold change)|>0 and adjusted p-values < 0.05 were considered to be differentially expressed and significant.

### Transcriptome Sequencing Analysis

Data processing and visualization were performed as previously mentioned.^40^ Briefly, raw data were uploaded to the Galaxy website (http://www.usegalaxy.org), quality checked (FastQC, Galaxy ver. 0.74) and mapped to the Mus musculus Genome Reference Consortium Mouse Build 39 (RNA Star, Galaxy ver. 2.7.11b). FeatureCount (Galaxy ver. 2.0.3) was used to count the number of reads per annotated gene in each mapped BAM file. Raw counts of genes were transferred to RStudio (ver. 2024.12.1) and normalized with the DESeq2 package (ver. 1.42.1). The heatmap was created with the heatmap package (ver. 1.0.12), and principal component analysis (PCA) plot and volcano plot were generated using the ggplot2 package (ver. 3.5.2). Functional annotation clustering using Gene Ontology (GO) and pathway analysis of DEGs with the Kyoto Encyclopedia of Genes and Genomes (KEGG) were conducted through the Database for Annotation, Visualization, and Integrated Discovery (DAVID) v2023q4.^41^

### Statistical Analysis

Statistical analysis was performed in GraphPad Prism. Data are expressed as mean ± S.E.M. Statistical analysis was performed using a one-way or two-way ANOVA with Tukey/Sidak’s/Dunnett’s multiple comparison test. *p ≤ 0.05, **p ≤ 0.01, ***p ≤ 0.001, ****p ≤ 0.0001.

## RESULTS AND DISCUSSION

### Self-assembly of KFE5

The assembly of the pentapeptide FKFEF in water was characterized using negative-stain TEM. Data shows that compared to KFE8 (FKFEFKFE), which assembles into thin nanofibrils (8-10 nm wide), the truncated variant KFE5 (FKFEF) exhibits markedly different assembly behavior, forming broader tape-like structures 50-100 nm wide (**Figure S9**). This significant increase in fibril width is attributed to a shift in the register of amino acid alignment during the assembly process. Notably, in KFE8, the terminal phenylalanine is out of register, which influences the stacking interactions and stabilizes the formation of narrow nanofibers.^42^ In contrast, the shorter KFE5 lacks this out-of-register terminal phenylalanine, leading to altered intermolecular interactions and a different registry of amino acids within the fibril core (**Figure S10**). This shift affects the packing density, favoring the formation of wider tapes rather than the narrower nanofibers observed in KFE8. To gain insights into KFE5 assembly, we employed molecular dynamics (MD) simulation of KFE5 based on an Alphafold3 prediction and the model developed for KFE8.^32^ The Alphafold3 predictions for 30 copies of KFE5 included one anti-parallel 2-start fiber, similar to the KFE8 model previously reported by us.^14,32^ In this model, significant interactions contributing to the self-assembly of the pentapeptide are the π-π stacking interactions between F-F in a hydrophobic core, sandwiched between the β-sheet-forming peptide backbone (**Figure 1A, B**). As described in the methods section, we extended this initial 30-peptide proto-fiber to a longer 192-peptide dual-fiber model for analysis using molecular dynamics. The KFE5 fiber has the basic configuration of the KFE8 and KFE12 peptide assemblies,^43^ featuring a hydrophobic core of phenylalanine sandwiched between the β-sheet-forming peptide backbone. One significant difference with KFE5 is the alignment of the phenylalanine residues along the edge of the fiber. This produces a hydrophobic strip along the edge of each fiber sandwich, creating a source for inter-fiber interactions. The alignment of the upper and lower sheets varies throughout the simulation, leaving an overlap at each edge, which could allow other fibers (**Figure 1C**) to join the assembly, which can now grow in both directions, along the fiber axis (z-plane) and in the plane of the β-sheet (x-plane). The edge interactions are almost exclusively hydrophobic and exclude solvent atoms from the fiber interface. These findings underscore the importance of peptide sequence length and amino acid register in modulating the morphology and dimensions of self-assembled peptide nanostructures.

**Figure 1.**
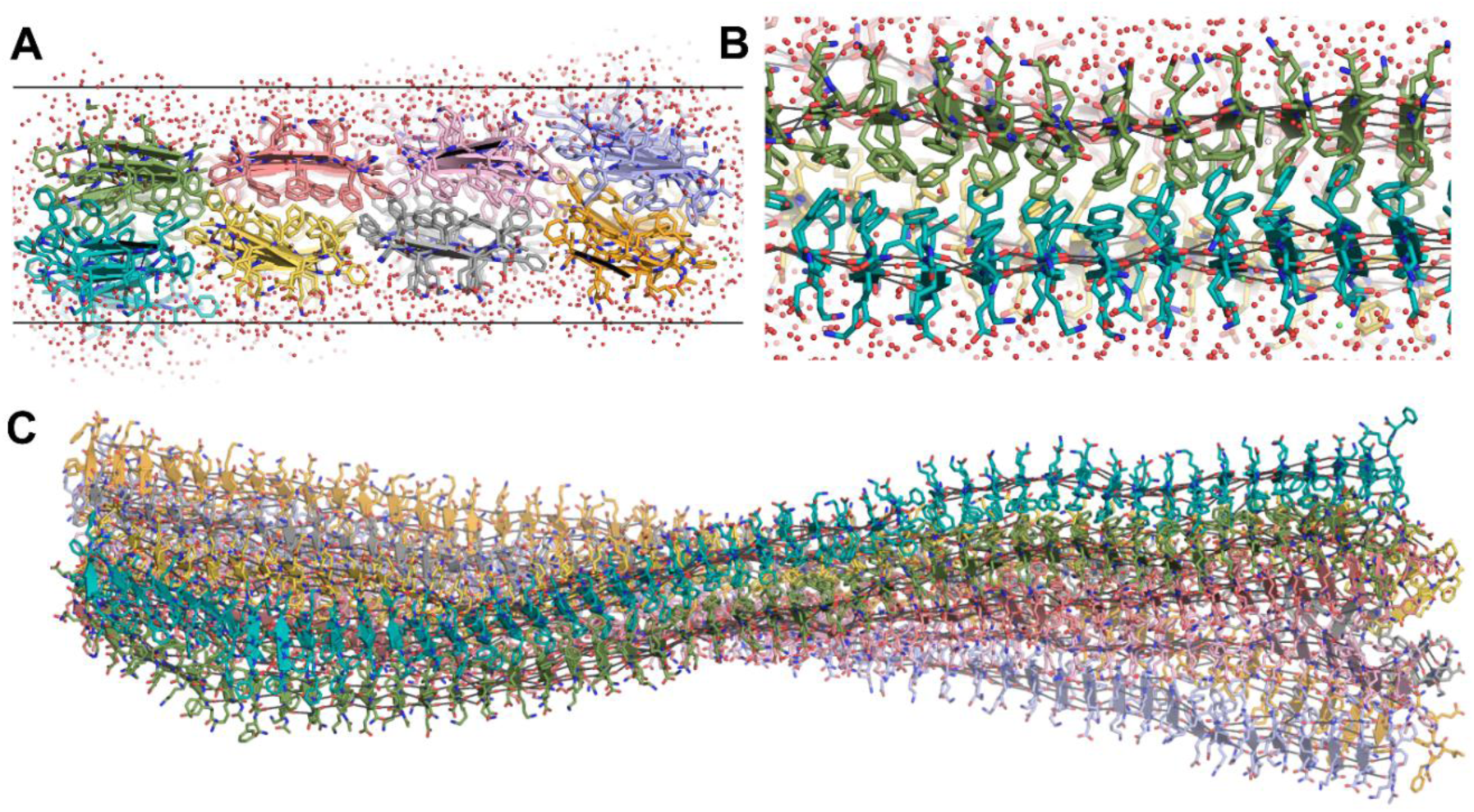
A model of the KFE5 assembly. (A) Close-up view of the KFE5 peptide assembly. The solvent layer around the peptide is shown. The two horizontal lines demark the circa 32 Å sheet thickness determined via SAXS. (B) Edge-on view of the assembly showing the phenylalanine hydrophobic core. The β-sheet forming hydrogen bonds are shown as lines. (C) The 384-peptide cartoon model of the two fibers interacting through their mixed hydrophobic and charged edges (Side-chains not shown for clarity).

### Substituent effects on the self-assembly of KFE5

In water and under identical preparation conditions, all peptides, except the hydroxyl (OH)-substituted variant, assembled into 3D nanostructures (**Figure 2A-H, S11**). Morphological differences among the fibrils were evident as the electronic properties of the substituents attached to the benzyl group modulate the strength of the π-π interactions, molecular packing, and the overall fibrillar architecture. EWGs decrease the negative charge density on the π-cloud, while EDGs exert the opposite effect via local dipole interactions, altering the aromatic quadrupole. In stark contrast with the unsubstituted parent peptide (X=H), which formed 50-100 nm wide tapes, the presence of EWGs, CN, and NO₂ resulted in fine and significantly thinner fibrils (∼15-17 nm wide). Interestingly, bromination resulted in dense, wide tapes with similar widths (∼40 nm wide) compared to the parent peptide KFE5.

**Figure 2.**
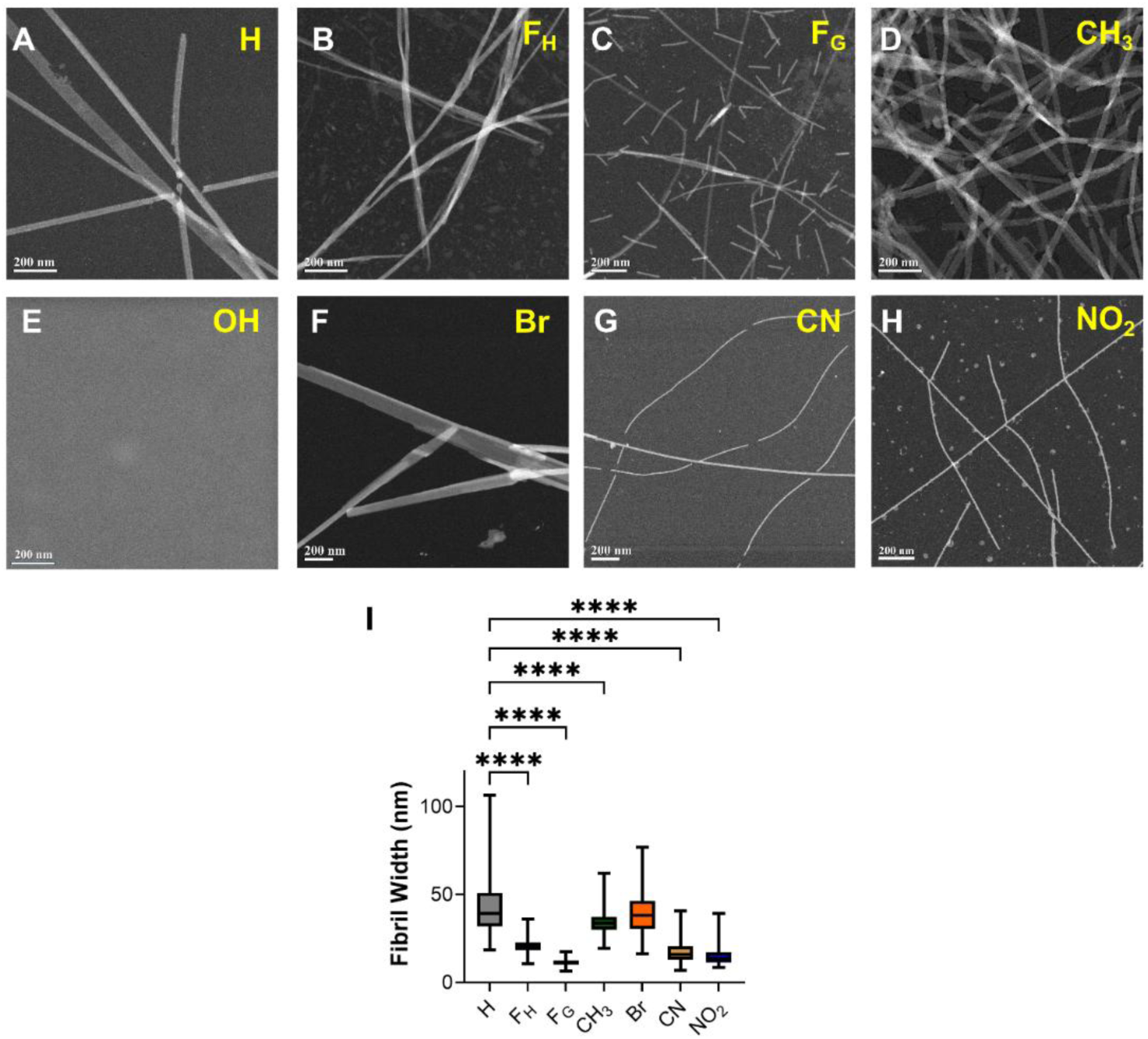
TEM images of self-assembled pentapeptides having different substitutions. (A) H, (B) F_H_, (C) F_G_, (D) CH_3_, (E) OH, (F) Br, (G) CN, (H) NO_2_ with difference in (I) fibril width as calculated from TEM images. ****p < 0.0001 as determined by a one-way ANOVA.

Substitution with the EDGs, OH, and CH₃ led to complete ablation or formation of dense twisted fibrils, respectively. These morphological diversities are likely due to the relative strengths of the EDGs, with the strong resonance active OH group inhibiting assembly and the weaker inductive electron donor CH_3_ promoting assembly, especially at the para position. The width of the methylated fibrils was comparable to the parent peptide (∼42 nm), and they were the only fibrils that exhibited a twisted morphology. Further, lengthening (F_H_) or shortening (F_G_) the benzyl group position from the peptide backbone also resulted in notable changes in fibril morphology. The F_G_ variant formed shorter fibrils, averaging 117±56 nm in length, whereas the F_H_ variant produced bundled thick fibrils with similar lengths compared to the KFE5. Interestingly, the fibril width was decreased for both F_H_ and F_G_ peptides, with the F_G_ peptides displaying the lowest width (15 nm). These differences probably arise from variations in side-chain volume and orientation within the hydrophobic bilayer, which influence peptide packing and assembly. The impact of these structural variations on fibril width is summarized in **Figure 2I**.

### Determination of secondary structure

Circular dichroism (CD) spectroscopy data indicate that KFE5 fibrils adopt a classical β-sheet signature, characterized by a negative peak centered around 220 nm (n–π* transition) and a positive peak near 198 nm (π–π* transition) (**Figure 3A**). Variations in the β-alkyl chain length modulated the spectral broadness; notably, the broad minima observed between 210-220 nm in the parent peptide shifted to sharper minima centered at ∼218 nm for both F_G_ and F_H_ (**Figure 3B, C**). The CH_3_-substituted analog retained the canonical β-sheet CD signature of the KFE5 peptide (**Figure 3D**). A strong positive CD signal was detected at ∼235 nm for the OH-substituted peptide (tyrosine) despite the lack of fibril formation by TEM (**Figure 3E**). This is a hallmark of tyrosine aromatic stacking interactions in peptide fibrils or aggregates, serving as a spectroscopic marker of such interactions.^44^ Substitutions with EWGs induced a shift in the CD spectra, with the brominated peptide displaying a negative ellipticity peak ∼240 nm (**Figure 3F**), attributable to electronic transitions associated with the amide groups and aromatic interactions within β-sheets. The CN and NO_2_ peptides, on the other hand, exhibited a positive Cotton effect at 240 nm, associated with π–π stacking and aromatic stacking interactions (**Figure 3G**, **3H**).

**Figure 3.**
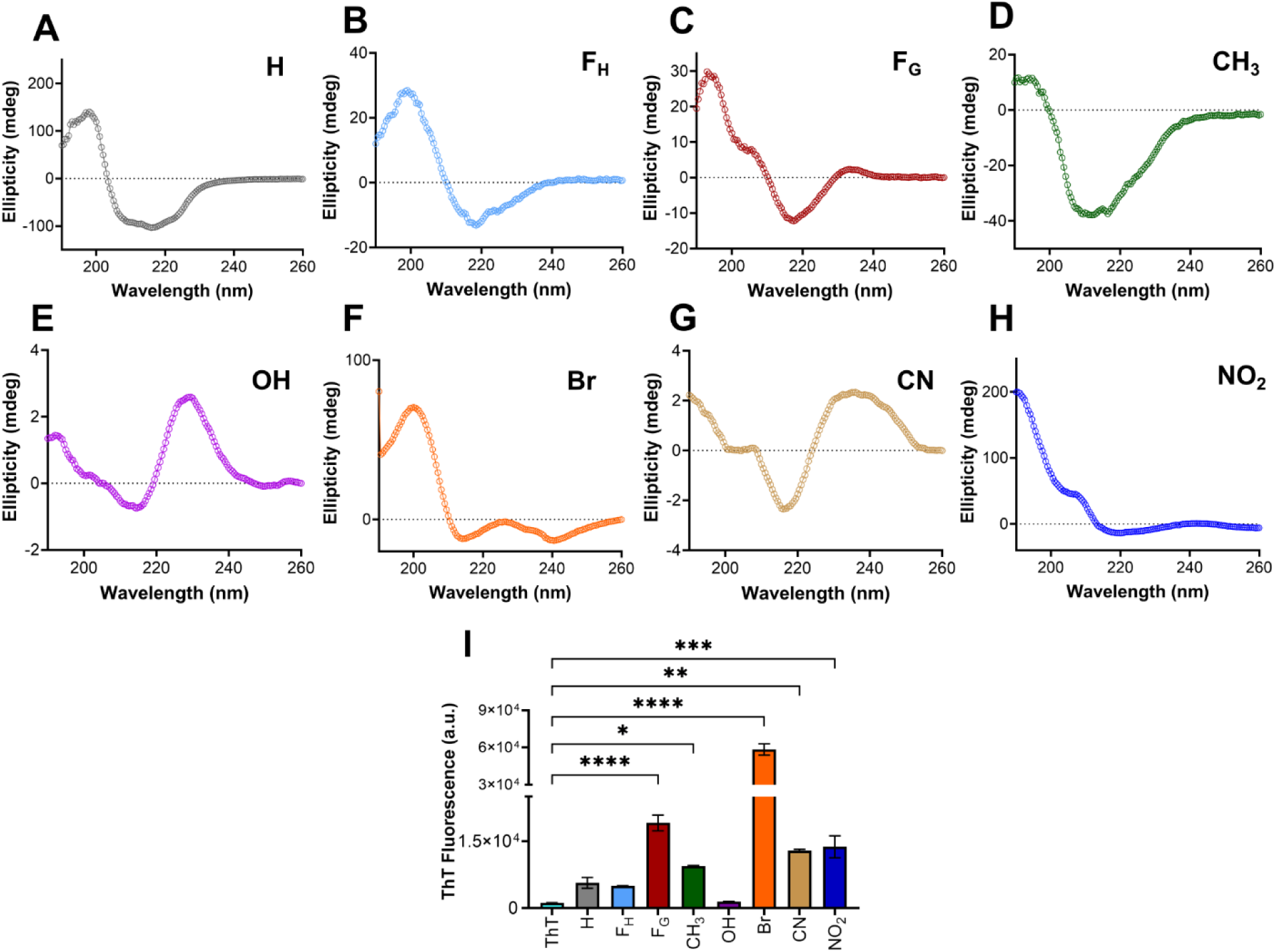
(A-H) CD spectra of self-assembled pentapeptides showing differences in secondary structures. (I) Variations in ThT fluorescence intensities in presence of different fibrils formed by pentapeptides as marked in the figure. *p < 0.05, **p < 0.01, ***p < 0.001, ****p < 0.0001 as determined by a one-way ANOVA.

To confirm the formation of β-sheet-rich fibrils, Thioflavin T (ThT) fluorescence assays were performed. ThT, a well-established amyloid fibril marker,^45–47^ exhibits increased fluorescence upon binding to β-sheet-rich structures due to restricted bond rotation. The brominated peptide produced the highest ThT fluorescence (**Figure 2I**), commensurate with the dense thick fibrils observed by TEM and β-sheet structure revealed by CD spectroscopy. The presence of EWGs CN and NO_2_ also led to an increased ThT signal, which was less than that of the strong halogen Br but higher than that of the parent peptide. Consistent with TEM data, the signal from the OH-substituted variant was the lowest and comparable to ThT alone, suggesting a lack of fibrils, whereas CH_3_ substitution promoted fibril formation and increased fluorescence. Interestingly, despite forming fibrils of lower length and width, fluorescence was enhanced for the F_G_ peptide compared to the parent peptide or the F_H_ peptide.

Complementing our CD data, FT-IR analysis of the substituted peptides revealed predominantly β-sheet characteristics, with major amide-I bands centered at 1623-1627 cm^-1^ (**Figure S12**). The additional peak, centered around 1679-1687 cm^-1,^ represents an anti-parallel β-sheet orientation in almost all the peptides. Fibrils formed from F_G_ peptide, however, do show a significant amount of disordered nature along with β-sheet characteristics, with a sharp peak at 1647 cm^-1^. Similarly, the anti-parallel β-sheet orientation with peaks centered around 1682 cm^-1^ is almost missing from the fibrillar pattern observed in the F_H_-substituted peptide, signifying the possibility of the formation of a parallel β-sheet in this case. Thus, the FT-IR analysis further confirms the β-sheet-rich secondary structure for all the fibrils formed by different pentapeptides.

### Substituent effects on viscoelastic properties of peptide hydrogels

The gelation propensity and viscoelastic behavior of peptide hydrogels with various substituents were characterized. Data indicated that despite robust assembly and a high β-sheet content, a 5 mM solution of the parent KFE5 peptide failed to form a solid gel (**Figure 4A**). Interestingly, all peptide variants bearing EWGs (Br, CN, NO_2_) formed self-supporting gels at the same concentration. In contrast, peptides substituted with EDGs (OH and CH_3_) also failed to undergo gelation at a peptide concentration of even 5 mM (**Figure 4A, S13**).^48^ All peptide solutions demonstrated viscoelastic properties, with the storage modulus (G′) exceeding the loss modulus (G″) and exhibited linear viscoelasticity up to 1% strain (**Figure 4B, C**). Notably, NO₂ substitution derivatives displayed the highest G′ values, whereas OH substitution resulted in a G′ approximately 2.5-fold lower than the parent peptide, as expected. Substitution with Br, CN, and CH₃ yielded intermediate G′ values.

**Figure 4.**
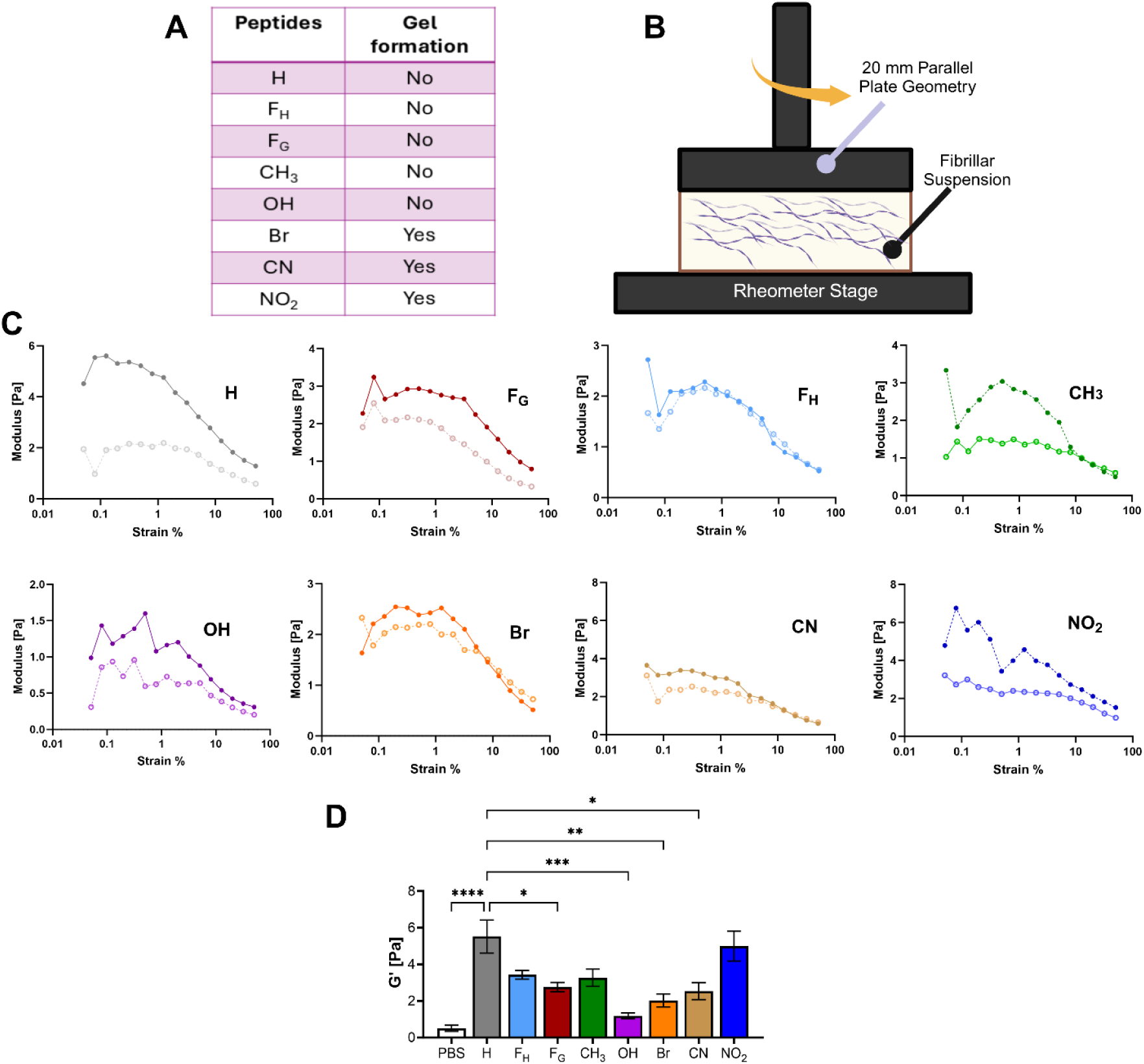
(A) Probability of gel formed by peptides with substitutions Br, CN, and NO_2_ at 5 mM peptide concentration. (B) Schematic of peptide nano-fiber solution testing using a rotational rheometer. (C) Representative plots showing the evolution of G′ (dark, solid circles) and G″ (light, open circles) as a function of strain amplitude. (D). *G*′ of peptide solutions at a strain of 1% and angular frequency of 10 rad/s. *p < 0.05, **p < 0.01, ***p < 0.001, ****p < 0.0001 as determined by a one-way ANOVA.

Complementary viscosity measurements corroborated these rheological trends, showing that NO₂ and CH₃ substitutions possessed the highest viscosities (**Figure 4D**), whereas Br and OH substitutions resulted in the lowest viscosities. These findings are supported by transmission electron microscopy (TEM) imaging, which revealed morphological differences consistent with the rheological data. The side chain length also influenced mechanical properties; peptides with an extended phenyl group (F_H_), formed thicker and longer fibrils, correlating with increased G′ and viscosity. These F_H_-hydrogels exhibited shear-thinning behavior, similar to that of F_G_ substitutions, indicating a potential for injectable applications. Overall, substituent-induced variations in fibril morphology directly impacted the viscoelastic properties of the peptide hydrogels, revealing the capacity to tune mechanical characteristics through side-chain chemistry.

### SAXS and WAXS analysis of the 5-mer peptide assemblies

All the SAXS curves fit with long polydisperse rods plus smaller components (**Figure 5A**). The characteristics of these rods, extracted from fitting to a Guinier-plate model, are almost identical with an average rod or tape thickness estimated at 31.9(1) Å (**Figure 5B**). This is similar to our previously measured values for KFE8 and KFE12.^14,43^ This thickness, which includes contributions from the solvent layer, matches the β-sheet sandwich model. This appears to be a common feature of the KFE peptides that is not disrupted by the modification of the phenylalanine side-chain, which forms an oily interface between the upper and lower β-sheets (**Figure 1A**). The width of these tapes, as measured by TEM (**Figure S11**), is at the limit of resolution of our SAXS data, about 60 nm (600 Å). Therefore, we cannot apply the standard Guinier radius of gyration, Guinier-Rod radial cross-section approximations, or Holtzer-plot analyses to the SAXS data, but the Guinier-plate analysis, which models an infinite sheet, is expected to be unaffected by this. The Debye-Bueche model of gel n-homogeneity is also likely to be unaffected. The normalized residuals of Debye-Bueche inhomogeneity-correlation length (ζ) plots (SAXS plots fitted to DB equation) have reasonable but systematic variations with minor errors in ζ, except for F_H_, indicating that the approximation is valid over the selected fitting range (0.015 - 0.10 Å^-1^). The gel inhomogeneity-correlation length (ζ) of the KFE5 peptides was similar (**Figure 5C**) in the range of 27 Å for NO_2_, 38-47 Å for CN, F_G_, and CH_3_, and about 58 Å for H and Br. The ζ of F_H_ was not well-determined, since the SAXS curve was not well-fit by the Debye-Bueche model. These compare with ζs for the KFE8 at 25 Å and KFE12 of 17 Å, depicting a trend towards larger ζ values for the shorter peptides.

**Figure 5.**
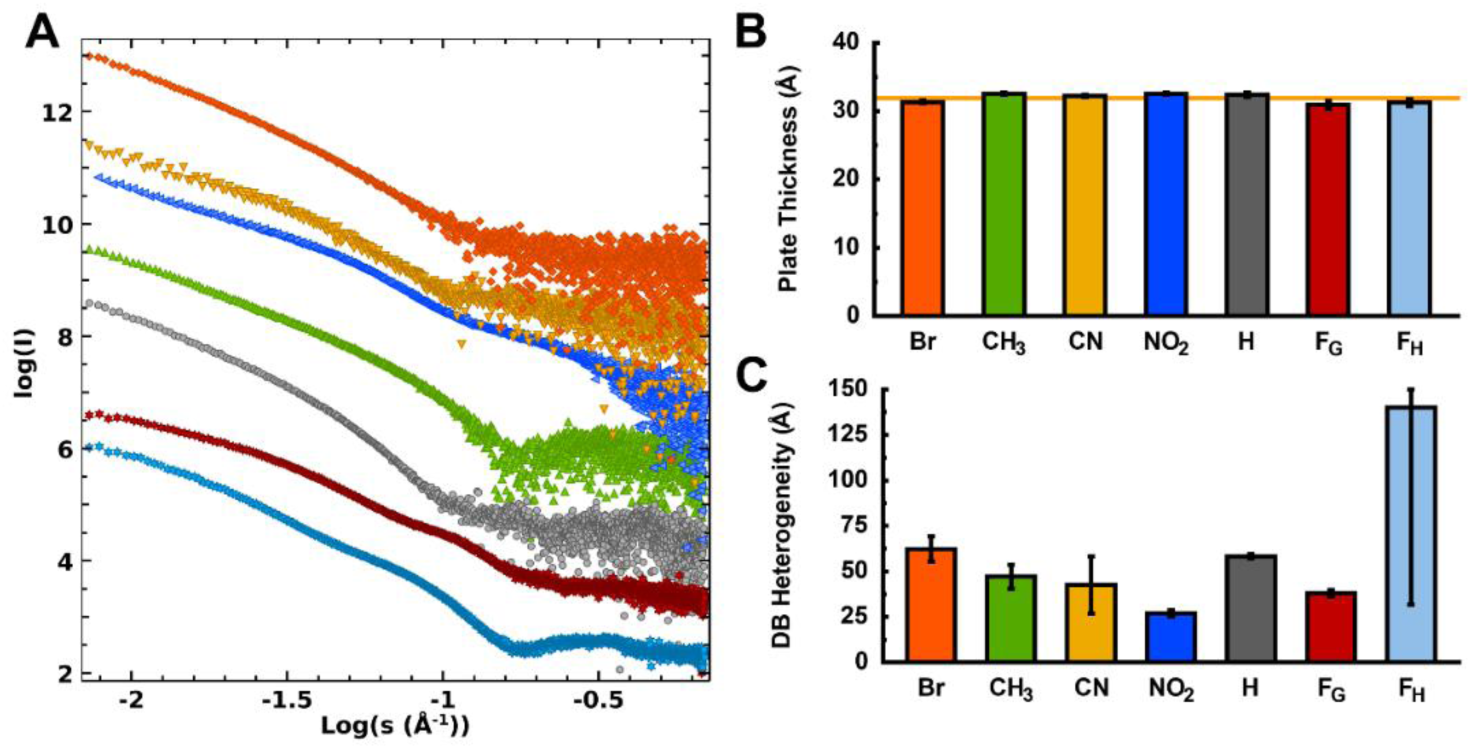
(A) The Log-Log 4 mg/mL SAXS curves for (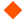) Br, (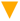) CN, (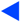) NO_2_, (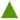) CH_3_, (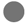) H, (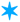) F_G_ and (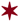) F_H_. SAXS curves are offset for clarity. (B) The GNOM Debye P(r) plate thickness fitting. The Debye P(r) was fit to a 6-term Fourier series with an exponential amplitude decay. The average Plate thickness, τ∼31.9(1) Å, is shown as an orange line. (C) The Debye-Bueche heterogeneity factors (ζ) for the KFE5 peptides. The value of ζ for F_H_ substituted peptide was undetermined.

X-ray diffraction scattering relies on a regular repeat of features in a molecular assembly and is an ideal tool for fibril-forming peptides such as KFE5. The β-sheet peak at d∼4.7 Å was the strongest in the X-ray diffraction scattering (XRD) for all KFE5 peptides (**Figure S14**). This phenomenon has also been observed in studies of other KFE peptides, specifically those of lengths 8 and 12, and is maintained in the KFE5 peptide fibers. The KFE5 peptide differs in the positioning of phenylalanine residues at both ends of the peptide, which modeling suggests aids in the formation of broad tapes formed by multiple fibers. Unique to KFE5, the second and third strongest XRD peaks are the d∼20 Å peak and the d∼24 Å peak from longer-range ordering of the fibers (**Figure S14, Table S1**). However, in the modified phenylalanine, these peaks are significantly weaker, dropping from ∼40% of the β-sheet peak intensity in KFE5 to 11% for CN, and 8% or lower for the other modified phenylalanine peptides. The shorter distance of 20 Å corresponds to the edge-to-edge fiber repeat distance (perpendicular to the fiber axis) in the KFE5 tape packing model. The weaker 24 Å peak matches the diagonal inter-fiber distance between the upper and lower β-sheets. The modified phenylalanine peptides have weaker edge fiber repeat peaks, indicating that they are less well ordered along the inter-fiber tape-forming direction.

### Effects of chemical substituents on innate immunity

Prior studies using KFE8 and other self-assembling peptide nanofibers have shown that they activate innate immune cells, such as dendritic cells (DCs) and macrophages, via the release of damage-associated molecular patterns (DAMPs).^49,50^ Studies have reported the upregulation of maturation markers, such as MHC II, CD80, and CD86, along with the secretion of chemokines (MCP-1α/CCL2, KC/CXCL1) and cytokines (GM-CSF, IL-5, IL-6, IL-1β), in response to treatment with β-sheet-rich peptide nanofibers.^51^ To address the effects of substituents on activation of innate immunity, we employed BMDC cultures and measured cytokine and chemokine release in response to treatment with the nanofiber variants. LPS (lipopolysaccharide), a component of bacterial cell walls and a potent activator of toll-like receptor 4 (TLR4), was used as a positive control. Analysis of the supernatant from treated BMDCs revealed that modifications to KFE5 have a significant influence on cytokine production and immunogenicity. Physical modification with F_G_ resulted in minimal cytokine release, characterized by extremely low levels of pro-inflammatory TNF-α, tolerogenic IL-10, and maturation-associated IL-13, along with negligible production of chemokines, including MCP-1, MIP-1α, MIP-1β, and IP-10. In contrast, F_H_ substitution led to the robust secretion of inflammatory cytokines, including IL-6, MCP-1, and TNF-α, along with moderate levels of IL-10 and MIP-1α **(Figure S15)**. Br substitution led to the secretion of IL-10 and MCP-1, IL-6, MIP-1α, and TNF-α compared to the parent peptide. Conversely, NO_2_ substitution did not induce IL-6 production, maintained MIP-1α levels, and specifically induced MIP-1β secretion. Of all the EWGs, the lowest activation profile was observed with CN substitution, with only moderate increases in MIP-1α and IL-13, similar to the unsubstituted peptide KFE5. Treatment with methylated fibrils resulted in the production of MIP-1α, MIP-1β, and TNF-α. In contrast, OH substitution resulted only in the release of MIP-1α and MIP-1β without TNF-α secretion, yielding a less immunogenic profile compared to CH_3_ substitution. These findings demonstrate that the presence of substituent groups on the benzyl ring can alter the innate immune activation profiles and could have potential implications for vaccine development.

### Substituent effects on amyloidogenesis and cross-seeding

Amyloid fibrils can spontaneously enter cells and seed the aggregation of various monomeric proteins, ultimately leading to cell death.^52^ Due to the structural similarity between synthetic peptide nanofibers and fibrils formed from pathological amyloids, we investigated whether substituents affect amyloidogenic propensity (**Figure 6**). We employed HEK293T α-synuclein biosensor cells, in which one copy of α-synuclein is fused to cyan fluorescent protein (CFP), while another copy is fused to yellow fluorescent protein (YFP). The CFP and YFP fused proteins, upon aggregation, can undergo fluorescence resonance energy transfer (FRET) due to their proximity. This is then quantified by measuring the FRET signal by flow cytometry (**Figure 6A**). We found that 50 nM concentration of α-synuclein pre-formed fibrils (PFFs) were able to cross-seed their monomeric form, as evident by the high percentage of cells with FRET positive signals (∼30%) (**Figure 6B, D**). In contrast, treatment with the peptide nanofibers did not lead to any detectable FRET signal (**Figure 6C, D**). These findings were corroborated using fluorescence microscopy, where abundant foci were observed upon incubating with α-syn PFFs, but not peptide nanofibers (**Figure 6C**). Interestingly, a 10-fold excess of the peptide concentration also did not enhance FRET signals, thereby further demonstrating the non-cross-seeding behavior of the peptide nanofibers (**Figure S16**). These data suggest that despite structural similarity, nanofibers formed via designed synthetic peptides and pathological amyloids are functionally distinct.

**Figure 6.**
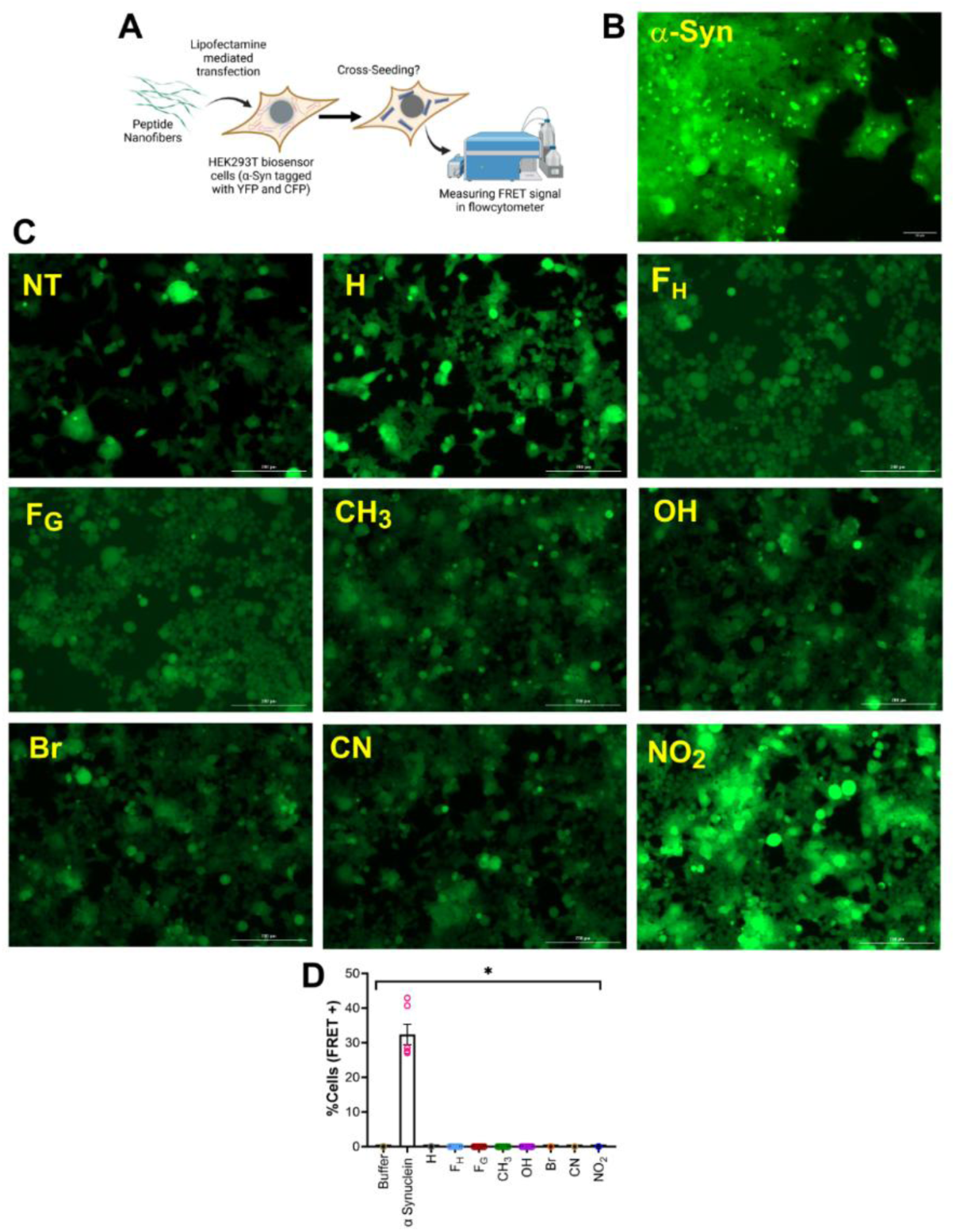
(A) Schematic representation of FRET assay in HEK293T biosensor cells in the presence of 5-mer peptide nanofibers. (B) Microscopy images of 50 nM α-Syn and (C) 5-mer PNFs added to biosensor cells. Scale bar 200 nm (D) % FRET positive cells as determined by the assay at 50 nM nanofiber concentration compared to α-synuclein PFFs (50 nM). *p < 0.05, **p < 0.01, ***p < 0.001, ****p < 0.0001 as determined by a one-way ANOVA.

### Immunogenicity of antigenic KFE5 variants

To assess the effects of substituents on the immunogenicity of the fibrils, we synthesized KFE5 variants in tandem with OVA, a peptide from chicken egg ovalbumin (aa 323-339 ISQAVHAAHAEINEAGR), which is an I-A^b^-restricted class II peptide (**Table S2, Figure S17-S24**). All OVA-bearing variants self-assembled into fibrils with diverse morphologies, which varied according to substitutions. The relative scarcity of fibril ends in TEM images suggested lengths on the order of microns, except for CN and F_G_ substitutions, which formed shorter fibrils (**Figure S25A**). The formation of β-sheet-rich fibrils was further confirmed by CD spectroscopy (**Figure S25B**).

Antigen presentation was quantified by using DOBW hybridoma cells, which produce IL-2 upon recognizing the OVA epitope in the context of MHC class II molecules on the surface of BMDCs (**Figure 7A**). The rationale underlying the method is to quantify IL-2 production, which is directly correlated to nanofiber internalization and processing by BMDCs. An advantage of this method compared to flow cytometry is that it eliminates false-positive readings resulting from fibril adhesion to the cell surface. Data indicated a concentration-dependent increase in IL-2 production in cultures treated with OVA-KFE5 variants (0.1-10 µM; 24h) (**Figure 7B**). Compared to the parent peptide, treatment with OVA-F_G_ variants produced higher levels of IL-2, presumably due to their shorter lengths that facilitate enhanced uptake. Differences in fibril morphology also affected internalization, where the thinner CN-substituted fibrils led to increased IL-2 production compared to the brominated variant, which aggregated into thick, dense aggregates. The other peptides showed moderate IL-2 production, which also increased with increasing peptide concentration (**Figure 7B**). We further evaluated persistence by treating BMDCs with 10 μM OVA-KFE5 variants and washing thoroughly before overlaying with DOBWs at different time points to remove extracellular fibrils (**Figure 7C**). Data indicated an increase in IL-2 production suggesover time for the parent peptide and F_G_ and F_H_ variants, indicating ongoing fibril processing and presentation in the context of MHC class II molecules. Except for fibrils formed by the parent peptide and F_G_ variant, IL-2 production decreased with time for all substitutions (**Figure 7C**). It is worth noting that these data also reflect the differences in internalization efficiency.

**Figure 7.**
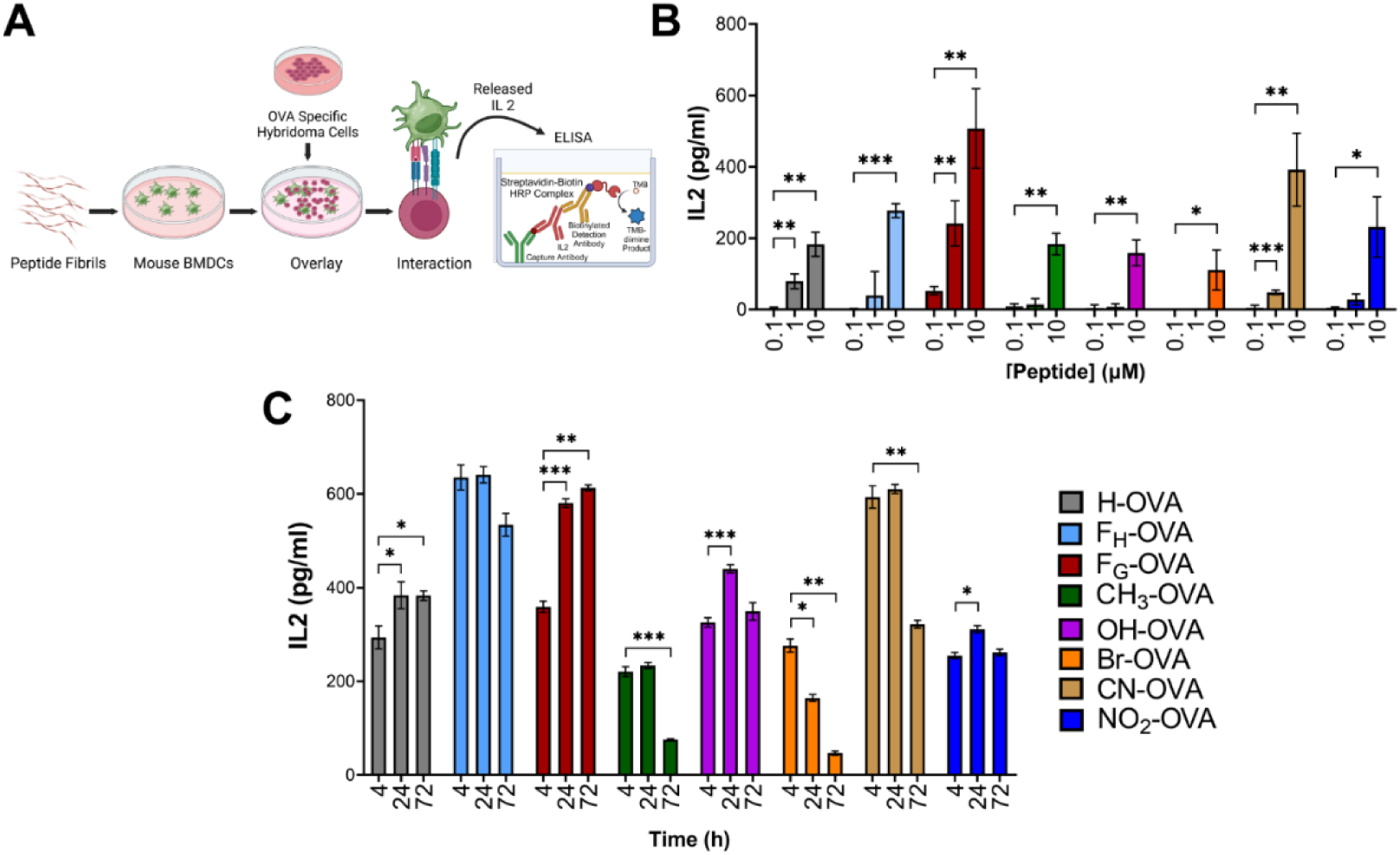
(A) Schematic representation depicting ELISA assay for IL-2 generation upon interaction of mouse BMDCs with DOBW hybridoma in the presence of 5-mer peptide nanofibers conjugated to OVA peptide. (B) Concentration dependent IL-2 production by OVA-conjugated different peptide nanofibers upon treatment in mouse BMDCs and overlayed with DOBW hybridoma for 16h. (C) Time dependent IL-2 production by OVA-conjugated different peptide nanofibers upon treatment in mouse BMDCs. p-values < 0.05 were considered statistically significant. *p < 0.05, **p < 0.01, ***p < 0.001, ****p < 0.0001.

### Antibody responses to chemically modified fibrils

To assess antibody responses, we primed and boosted groups of C57BL/6 mice with the OVA-bearing KFE5 variants and assayed the levels of anti-OVA antibodies in the sera (**Figure 8A**). Data indicated that, similar to KFE8, KFE5 acts as an immune adjuvant, leading to the production of anti-OVA antibodies (**Figure 8B**). Antibody responses were higher in mice vaccinated with OVA-bearing F_H_ and F_G_ variants, and this data agrees with our *in vitro* antigen presentation assays (**Figure 7**). Interestingly, a stark contrast in immunogenicity was observed for the EDGs with an OH substitution, resulting in robust antibody production, which was completely abrogated when a CH_3_ group replaced the OH group (**Figure 8B**). A similar effect was observed with the EWGs, where NO_2_-KFE5 fibrils bearing OVA were not immunogenic compared to Br or CN-substituted fibrils that resulted in robust antibody responses (**Figure 8B**). Analysis of the antibody isotypes revealed that, similar to KFE8, KFE5, and F_H_ peptides, these induced a Th2-biased response with IgG1 but no detectable IgG2c. In contrast, IgG1 and IgG2c were detected in the sera of mice vaccinated with the OVA-F_G_ variant, indicating a Th1/Th2 balanced response (**Figure 8C**). Similar to KFE5, OH modification resulted in IgG1 production alone, whereas brominated and CN-substituted fibrils induced both IgG1 and IgG2c antibody isotypes (**Figure 8C**). We next analyzed the tetramer^+^ CD4^+^T cell populations following splenic antigen recall in the spleens of vaccinated mice. Data show the presence of antigen-specific CD4^+^ T cells in almost all groups. A higher percentage of cells were detected in mice immunized with F_G_-OVA fibrils compared to other groups or controls (**Figure S26**). These findings are exciting because they show that substituents can not only influence the strength of the immune response but also its composition, opening up possibilities for tuning the immune response to the desired correlations of protection.

**Figure 8.**
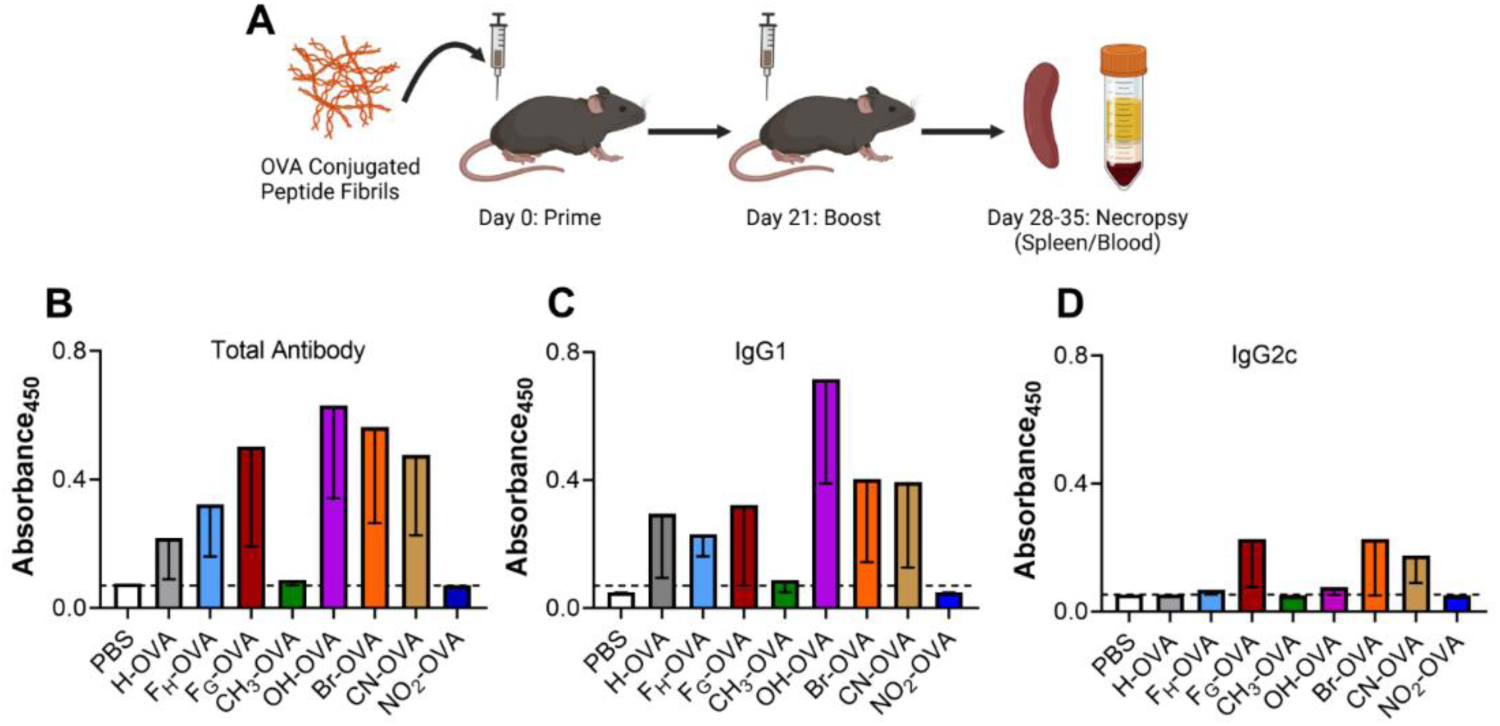
(A) Mice were primed with a subcutaneous dose of 100 μL of 2 mM peptide nanofibers and then given a subcutaneous boost of 2 mM PNFs 3 weeks later. At 1-2 weeks post-boost, blood sera were analyzed for antibody production (n = 6). (B) Total antibody production in vaccinated mice along with (C) IgG1 and (D) IgG2c isotypes.

### Transcriptomic sequencing analysis

To investigate how structural modifications of FKFEF peptides impact the transcriptomic profiles of BMDCs, we performed RNA sequencing on cells treated with various peptide constructs as shown in **Figure 9B**. Samples within each treatment group clustered tightly, indicating high intra-group consistency. All peptide-treated groups were distinctly separated from the non-treated (NT) group along PC1. Constructs modified with CN, NO₂, and OH groups closely overlapped with FKFEF, whereas those with Br and CH₃ substitutions were more distinct from FKFEF along PC1. Additionally, the separation along PC2 for (F_G_, H, and F_H_) suggested that changing the residual chain length contributed to further transcriptomic divergence not captured by PC1. These trends were corroborated by heatmap (**Figure 9A**) and volcano plots (**Figure 9C**), which highlighted consistent patterns of gene regulation. The FKFEF peptide significantly upregulated 172 genes and downregulated 179 genes. Further functionally annotating these differentially expressed genes (DEG) with GO and KEGG, we found that upregulated DEGs stimulated by FKFEF peptide were significantly enriched in biological processes such as the collagen fibril organization, antigen processing and presentation via MHC class I pathway, and positive regulation of T cell-mediated cytotoxicity (**Figure 9D**), as well as natural killer cell mediated cytotoxicity pathways (**Supplementary Table**). Compared to FKFEF peptide, the Br-substituted variant induced DEGs associated with positive regulations of ERK1 and ERK2 cascade, gene expression, and cell migration. In contrast, the downregulated DEGs from CH_3_ and F_G_ modification were involved in the IL-17 and TNF signaling pathways. The unregulated DEGs from F_H_ modification also increased the positive regulations of gene expression and MAP kinase activity (**Supplementary Table**).

**Figure 9.**
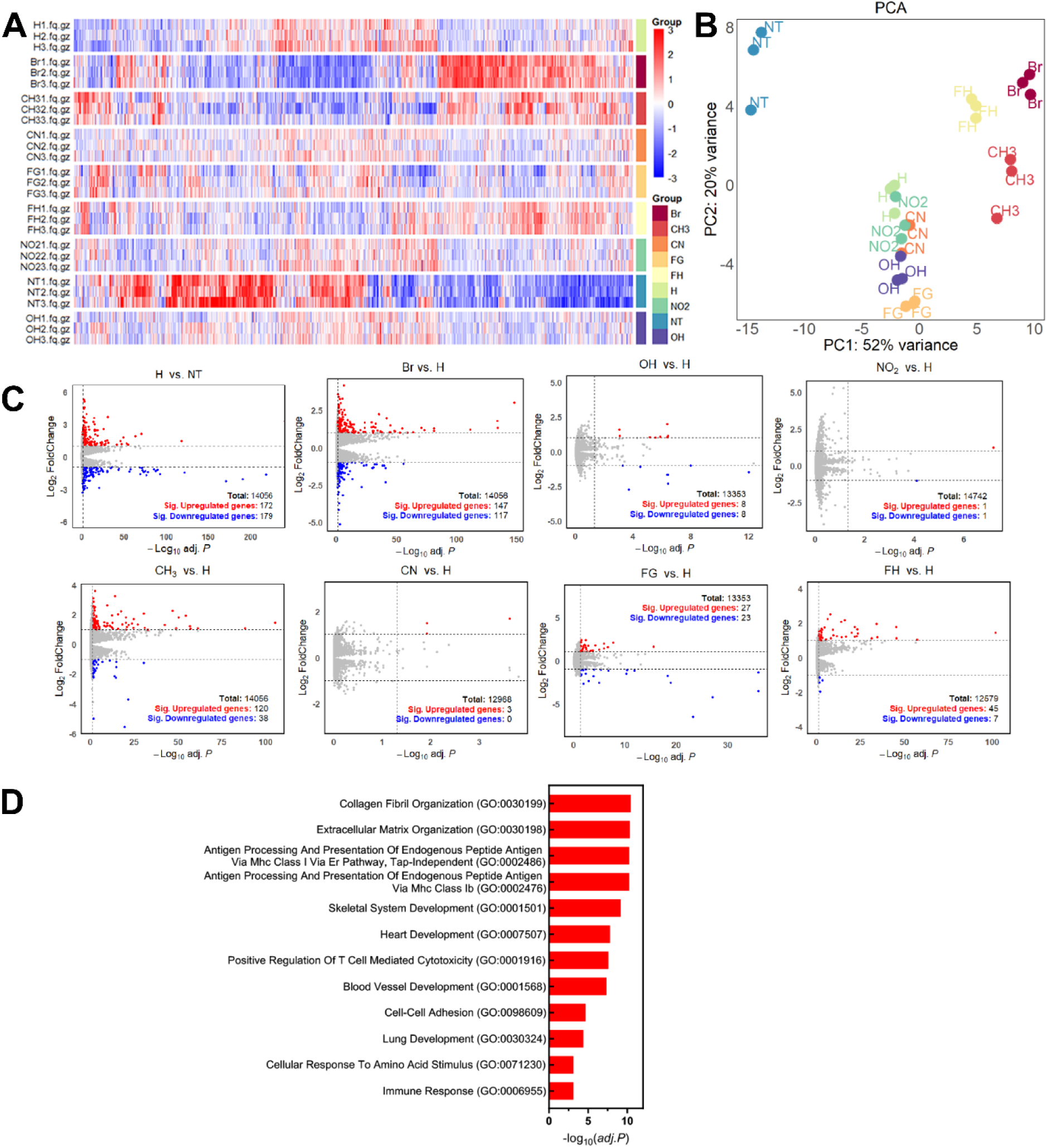
Transcriptomic sequencing analysis of mouse BMDCs treated with various kinds of non-natural amino acid peptides. (A) Heatmap showing the normalized counts of all differentially expressed (*adj.* p< 0.05) gene clusters of each peptide group compared with the FKFEF peptide. Each count in the gene column was scaled using Z-score across all samples. (B) PCA of mRNA expression variations in BMDCs treated with different peptides. (C) Volcano plots comparing gene expression levels between each modified peptide group to the parent peptide. Genes with significantly high expressions are shown in red (*adj.* p<0.05 and log_2_ fold change>1), while genes with significantly low expression (*adj.* p<0.05 and log_2_ fold change<-1) are shown in blue. Gray denotes genes that do not exhibit significant differences (*n*=3). (D) GO enrichment of unregulated DEGs after FKFEF peptide treatment.

## DISCUSSION

Supramolecular peptide assemblies are versatile nanomaterials used in vaccine development and immunotherapy, due to their self-adjuvanting properties.^7,53–56^ Previous research has enhanced immune responses by modifying charge, length, chirality, hydrophobicity, and secondary structure through chemical changes or the use of non-natural amino acids.^54,57–59^ Studies on sequence patterning, length, and substituent effects on (FKFE) repeats have deepened understanding of molecular packing and assembly modes.^4,19^ Building on this foundation, we demonstrate that KFE5 is the shortest amphipathic peptide capable of self-assembly. TEM analyses reveal that removing three amino acids from KFE8 to KFE5 causes a significant morphological change from narrow, nanofibrillar structures (∼8-10 nm wide) to broader, tape-like formations (50-100 nm wide). Molecular dynamics simulations suggest that a shift in phenylalanine stacking interactions accounts for this transition. In KFE8, an out-of-register phenylalanine at the terminus stabilizes narrow fibrils via specific π-π stacking, forming dense, uniform nanofibrils. Without this out-of-register phenylalanine in KFE5, stacking patterns change, promoting lateral association and wider tape formation. Simulations also show that phenylalanine residues in KFE5 tend to align along fiber edges, creating hydrophobic strips that support inter-fiber interactions and lateral growth. This edge hydrophobicity likely allows fibers to extend both longitudinally and laterally, explaining the broader morphology seen experimentally. The dynamic β-sheet alignment observed in the simulations suggests that edge interactions function as nucleation points for further fiber association, leading to variability in fibril width and increased lateral bundling.

Chemical modifications to the aromatic ring have a profound influence on fibril architecture. Electron-withdrawing groups (EWGs) such as CN and NO₂ produce markedly thinner fibrils compared to KFE5 tapes, indicating that EWGs weaken π-π stacking by reducing electron density.^60^ These modifications likely favor more planar, less densely packed arrangements, resulting in thinner, more flexible fibrils. Bromination, despite being an EWG, results in broader and denser tapes, suggesting steric effects and the specific electronic nature of the substituent modulate assembly differently than simple electron-withdrawing effects.^61,62^ Electron-donating groups (EDGs), such as OH and CH₃, exhibit contrasting effects: the hydroxyl group impedes fibril formation altogether, possibly due to excessive hydrogen bonding or disruption of aromatic stacking, whereas CH_3_ promotes fibril assembly but results in twisted fibrils.^63,64^ The twisting is presumably due to side-chain volume and polarity, which can induce conformational strain during the assembly process. Also, positional modifications of the benzyl group, such as lengthening (F_H_) or shortening (F_G_), further modulate fibril dimensions. The F_G_ variant produces shorter and thinner fibrils, likely due to altered packing geometry, whereas the F_H_ variant forms bundled, thick fibrils similar in length to KFE5, suggesting that increased side-chain length enhances lateral association through hydrophobic bridging.^65^

Spectroscopic data indicate that most variants adopt canonical β-sheet conformations, with subtle differences in spectral broadness and minima reflecting variations in packing and secondary structure. Notably, the OH-substituted peptide shows a strong positive CD signal at ∼235 nm despite the absence of visible fibrils in TEM, suggesting that aromatic stacking can occur independently of fibril formation.^66^ Thioflavin T fluorescence assays further confirm the presence of β-sheet-rich fibrils, with the highest fluorescence observed in the brominated peptide correlating with dense, thick fibrils seen by TEM. Interestingly, peptides with EWGs produce higher ThT signals, indicating enhanced β-sheet stabilization or density, while the hydroxyl variant shows low signals, consistent with its inability to form fibrils.^67^ FT-IR spectra support these findings, confirming the predominance of β-sheet secondary structures. Subtle spectral variations imply differences in β-sheet orientation, with the F_H_ variant favoring parallel β-sheets.^68^ These structural and morphological differences, resulting from the side chain chemistry, influence gelation and mechanical properties. While KFE5 alone does not gel at 5 mM, all EWG-containing derivatives (Br, CN, NO_2_) successfully form self-supporting gels, suggesting that electronic effects stabilize network formation through enhanced fibril interactions. Conversely, EDG substitutions (OH and CH_3_) hinder gelation, indicating that fibrillization alone is insufficient for gel formation and that network connectivity and side-chain interactions are crucial. The NO_2_ and CH_3_ derivatives exhibit higher storage moduli (G′) and viscosities, indicating mechanical robustness that can be tailored for biomedical applications. Prior studies have investigated the effects of different halogens on various positions of the benzyl ring using short hexamer peptides, demonstrating that increasing the degree of halogenation enhances the strength of stacking interactions and that heavier halides lead to stronger interactions.^31,69,70^ Despite structural similarities to pathogenic amyloids, modified KFE5 peptide fibrils do not promote cross-seeding of α-synuclein aggregation.

A key finding in this study is that fibril chemistry influences both innate and adaptive immune responses. Primary BMDCs treated with fibrils containing EWGs (Br, NO_2_, and CN) triggered a pro-inflammatory reaction compared to those treated with fibrils containing EDGs (OH, CH_3_). The F_H_ variant also induced a robust proinflammatory cytokine profile (e.g., IL-6, TNF-α) compared to the F_G_ variant, which elicits minimal cytokine secretion, suggesting that fibril density and surface chemistry modulate dendritic cell activation. These findings agree with immunological studies using other organic and inorganic nanomaterials.^71,72^ Variants like F_G_, with shorter fibrils, elicited higher IL-2 production from BMDCs, indicating efficient antigen internalization and processing. This effect was also observed in chemically modified fibrils, where antigen presentation capacity increased as fibril thickness decreased (CN>NO_2_>Br). In vivo studies demonstrated that KFE5 was also self-adjuvanting, similar to KFE8.^7^ The adjuvanting potency of positional substituents F_H_ and F_G_ was higher, eliciting stronger antibody responses with shorter F_G_ fibrils, which promoted a balanced Th1/Th2 profile, as indicated by the distribution of IgG isotypes. This improved adjuvant potency was also reflected in the percentage of OVA-specific CD4^+^ T cells in the spleens of vaccinated mice (**Figure S26**). Amongst the EDG substituted peptides, CH_3_ fibrils failed to elicit anti-OVA antibodies, whereas the OH substitution resulted in a Th2 biased response characterized by robust IgG1 production. In peptides bearing EWGs, no antibody production was detected for the NO_2_ variant, whereas Br and CN substitutions led to a balanced Th1/Th2 response with detectable IgG1 and IgG2c. Interestingly, despite the lack of robust antibody production, antigen-specific CD4^+^ T cells were detected in mice vaccinated with CH_3_ and NO_2_ fibrils, indicating that antigen processing and presentation to T cells occurred (**Figure S26**). Additional studies will be necessary to elucidate the immunological mechanisms underlying these findings.

The RNA-seq analysis reveals that peptides with CN, NO₂, and OH groups exhibit gene expression profiles similar to KFE5. In contrast, substitutions such as Br and CH₃ result in distinct transcriptomic shifts, highlighting the role of specific chemical groups. Additionally, modifications influencing residual chain length (F_H_) further diversify gene expression, highlighting the importance of peptide backbone structure. Functionally, KFE5 primarily upregulates genes related to immune processes, including collagen organization, antigen presentation, and cytotoxicity, highlighting its immunomodulatory potential. Variants like Br and differentially regulate pathways including ERK signaling, gene expression, and inflammatory responses, indicating that peptide modifications can produce tailored effects. These results demonstrate that peptide structure can modulate immune-related gene networks in DCs, with significant implications for the design of targeted therapies and immunotherapeutics. For instance, incorporating EWGs such as CN or Br enhances immunogenicity and promotes a balanced Th1/Th2 response, which is desirable for many protective vaccines. Conversely, modifications like F_G_ and OH can be tailored to favor specific immune profiles or minimize inflammation.

Overall, our data suggest that the differential immunogenic profiles driven by substituents demonstrate that chemical tuning offers a versatile approach to optimize peptide nanofiber vaccines. This level of control enables the design of personalized or disease-specific immunotherapies, where the nature and strength of immune responses are critical.

## CONCLUSION

In summary, our results demonstrate that chemical substituents on KFE5 nanofibers serve as potent regulators of their immunological and structural properties. By modifying side-chain chemistry, it is possible to improve fibril shape, influence innate immune activation, and direct adaptive immune responses. This research advances the intentional design of peptide-based nanomaterials for immunotherapy, vaccination, and other biomedical applications, emphasizing the essential role of chemical context in shaping biological outcomes.

## Supporting information

Table for Supporting Information

## ACKNOWLEDGEMENTS

Research funding was provided by the Washington University McKelvey School of Engineering, Department of Biomedical Engineering Commitment Funds (12–360–94361 J), and the National Institute of Allergy and Infectious Diseases (R01 AI168918) to Dr. Jai Rudra. Dr. Meredith Jackrel acknowledges support from grant R35GM153303 for research funding. The authors acknowledge the Sealy Center for Structural Biology and Molecular Biophysics at the University of Texas Medical Branch at Galveston for providing research resources. DOBW cells were a kind gift from Dr. Clifford V. Harding at Case Western Reserve University, Cleveland, OH, USA.

## AUTHOR CONTRIBUTIONS

Conceptualization, A.D., U.P, and J.S.R.; Methodology, C.B., S.P.Z., M.E.J., M.A.W., J.S.R.; Investigation, A.D., U.P., E.M.B., C.Y.L., H.G., J.F., M.G., S.S.; Writing – Original Draft, A.D., U.P, and J.S.R.; Writing – Review & Editing, C.B., S.P.Z., M.E.J., M.A.W., J.S.R.; Funding Acquisition, M.E.J., J.S.R.; Resources, C.B., S.P.Z., M.E.J., M.A.W., J.S.R.; Supervision, J.S.R.

## DATA AVAILABILITY

Sequence raw data files are available at NCBI BioProject accession #PRJNA1283513. Other data will be made available upon request.

