## Supporting Information for "Substituent-based Modulation of Self-Assembly and Immunogenicity of Amphipathic Peptides"

**Table of Contents**

**Figure S1**. MALDI-TOF and HPLC spectra of KFE5	S3

**Figure S2**. MALDI-TOF and HPLC spectra of KFE5(F_H_)	S4

**Figure S3**. MALDI-TOF and HPLC spectra of KFE5(F_G_)	S5

**Figure S4**. MALDI-TOF and HPLC spectra of KFE5(CH_3_)	S6

**Figure S5**. MALDI-TOF and HPLC spectra of KFE5(OH)	S7

**Figure S6**. MALDI-TOF and HPLC spectra of KFE5(Br)	S8

**Figure S7**. MALDI-TOF and HPLC spectra of KFE5(CN)	S9

**Figure S8**. MALDI-TOF and HPLC spectra of KFE5(NO_2_)	S10

**Figure S9.** TEM images of KFE8 and KFE5	S11

**Figure S10.** A 500 ns MD Simulation of the AlphaFold3 KFE5 model	S12

**Figure S11**. TEM images for KFE5 variants	S13

**Figure S12.** FT-IR spectroscopy of KFE5 variants...	S14

**Figure S13.** Representative images of gel formation	S15

**Figure S14.** Powder diffraction peak fitting for KFE5 variants	S16-S17

**Table S1**. WAXS peaks for KFE5 peptides	S18

**Figure S15**. Cytokine and chemokine production by KFE5 variants	S19

**Figure S16.** FRET analysis for α-synuclein aggregation by KFE5 variants (0.5 µM)	S20

**Table S2.** Sequences for OVA conjugated KFE5 variants	S21

**Figure S17.** MALDI-TOF and HPLC spectra of KFE5-OVA	S22

**Figure S18.** MALDI-TOF and HPLC spectra of KFE5(F_H_)-OVA	S23

**Figure S19.** MALDI-TOF and HPLC spectra of KFE5(F_G_)-OVA	S24

**Figure S20.** MALDI-TOF and HPLC spectra of KFE5(CH_3_)-OVA	S25

**Figure S21.** MALDI-TOF and HPLC spectra of KFE5(OH)-OVA	S26

**Figure S22.** MALDI-TOF and HPLC spectra of KFE5(Br)-OVA	S27

**Figure S23.** MALDI-TOF and HPLC spectra of KFE5(CN)-OVA	S28

**Figure S24.** MALDI-TOF and HPLC spectra of KFE5(NO_2_)-OVA	S29

**Figure S25.** TEM and CD of OVA conjugated KFE5 variants	S30

**Figure S26.** Percent OVA-specific CD4^+^T cells	S31

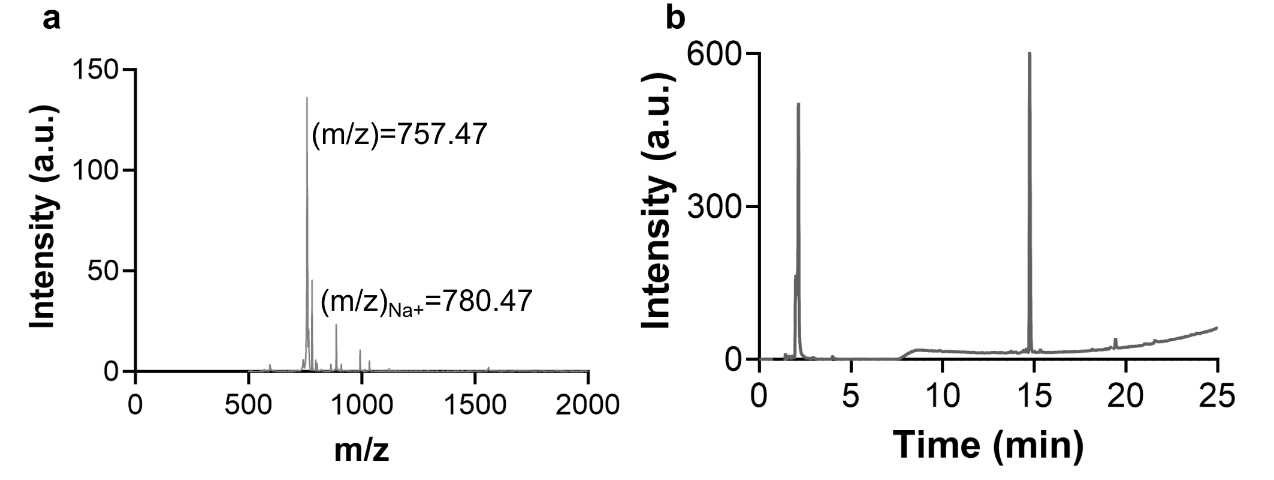

**Figure S1.** (a) MALDI-TOF and (b) HPLC profiles of KFE5.

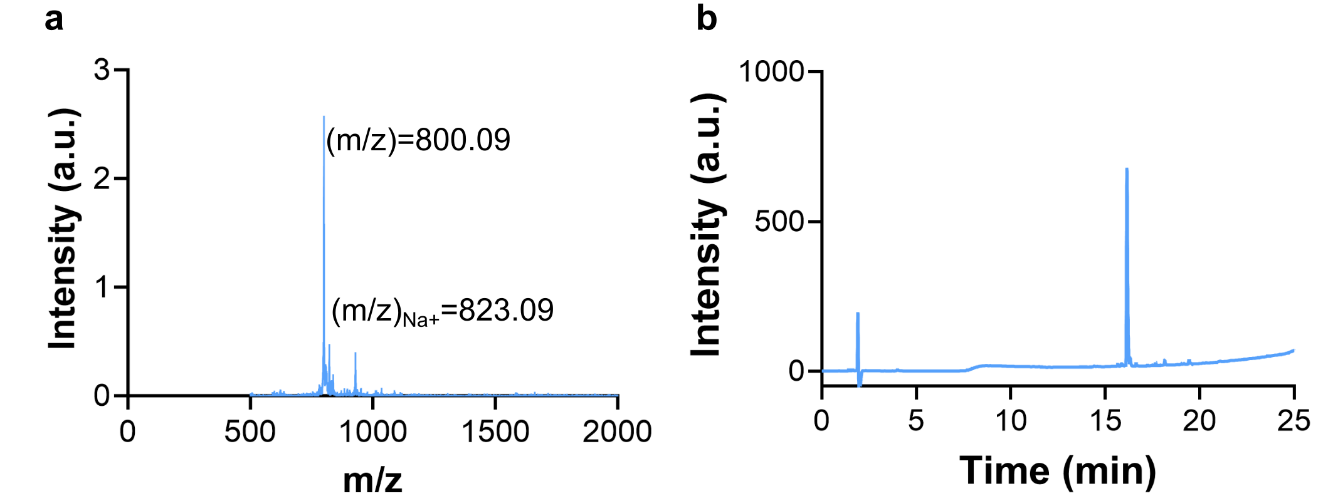

**Figure S2.** (a) MALDI-TOF and (b) HPLC profiles of KFE5(F_H_).

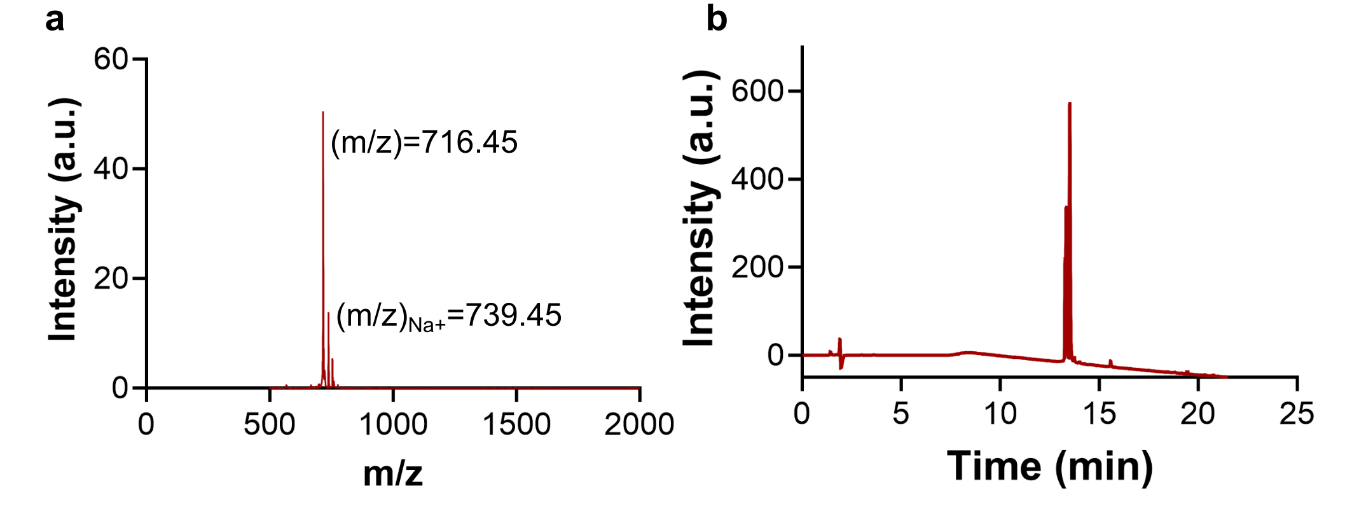

**Figure S3.** (a) MALDI-TOF and (b) HPLC profiles of KFE5(F_G_).

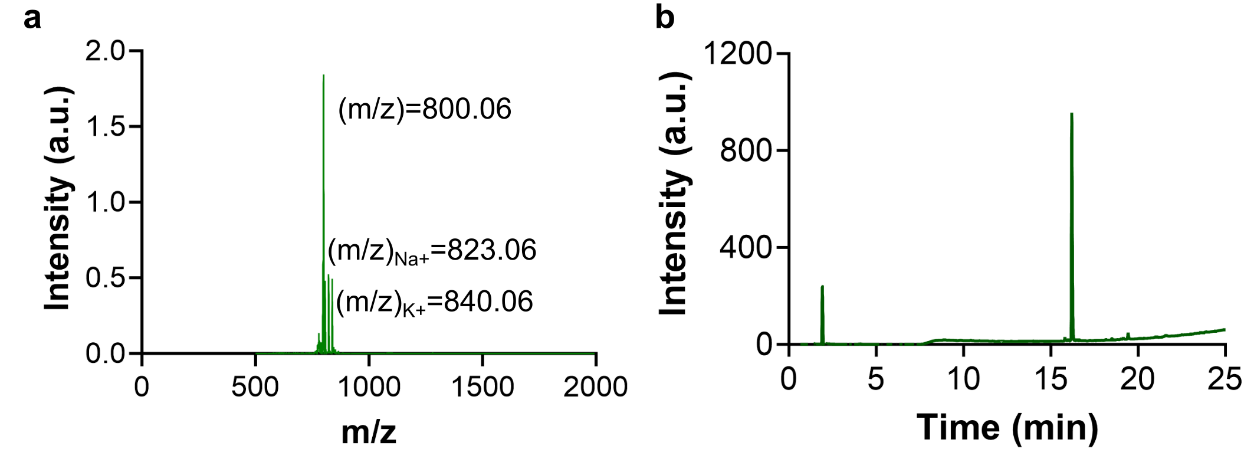

**Figure S4.** (a) MALDI-TOF and (b) HPLC profiles of KFE5(CH_3_).

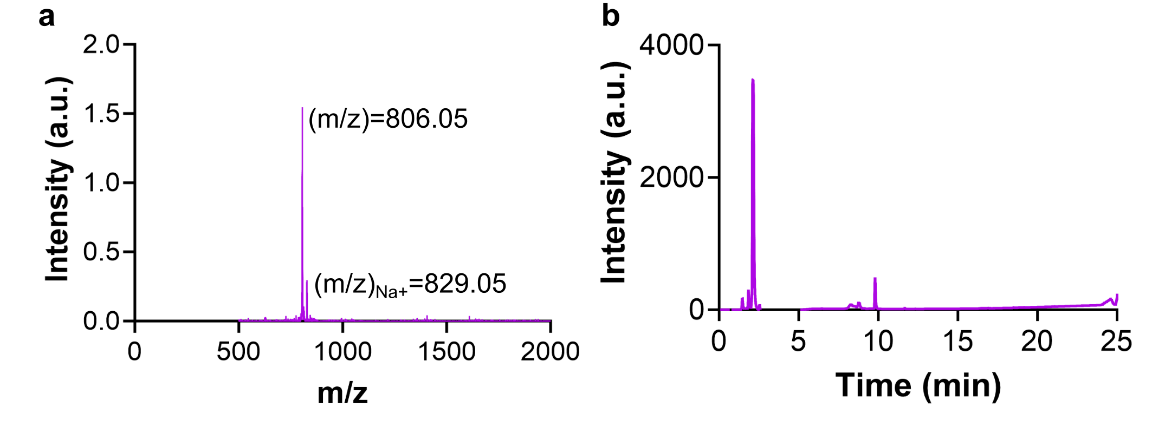

**Figure S5.** (a) MALDI-TOF and (b) HPLC profiles of KFE5(OH).

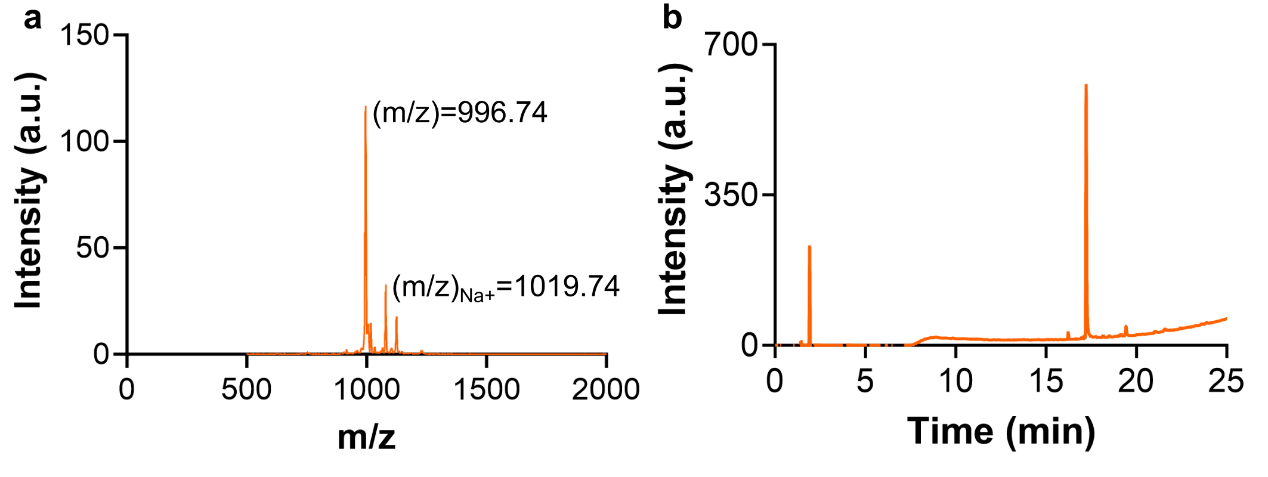
**Figure S6.** (a) MALDI-TOF and (b) HPLC profiles of KFE5(Br).

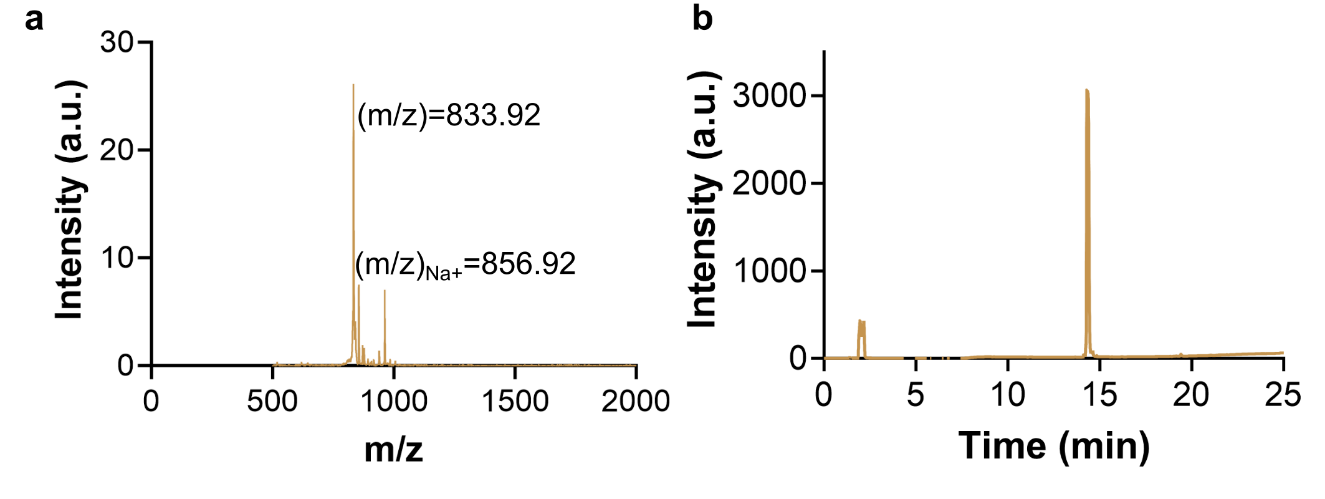

**Figure S7.** (a) MALDI-TOF and (b) HPLC profiles of KFE5(CN).

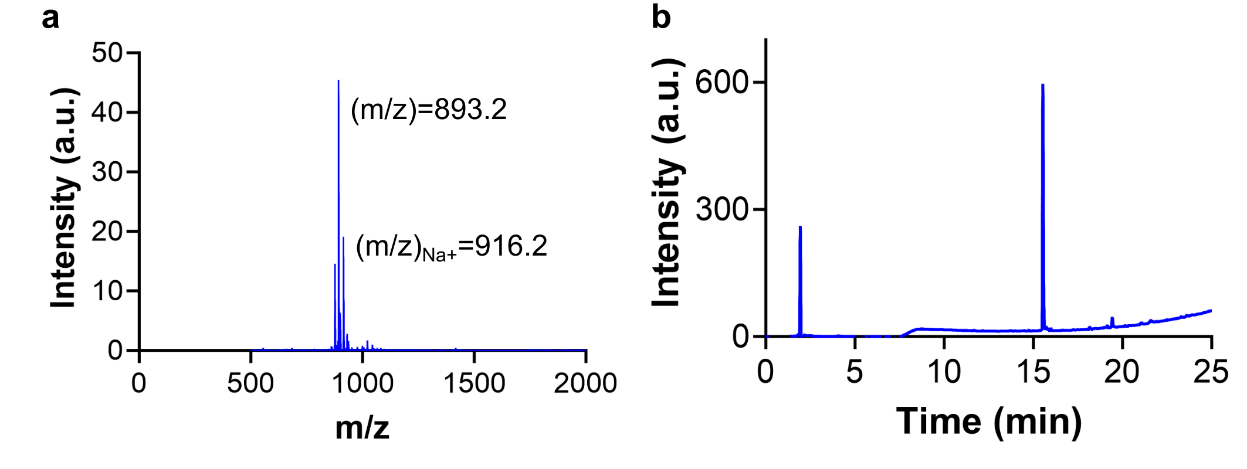

**Figure S8.** (a) MALDI-TOF and (b) HPLC profiles of KFE5(NO_2_).

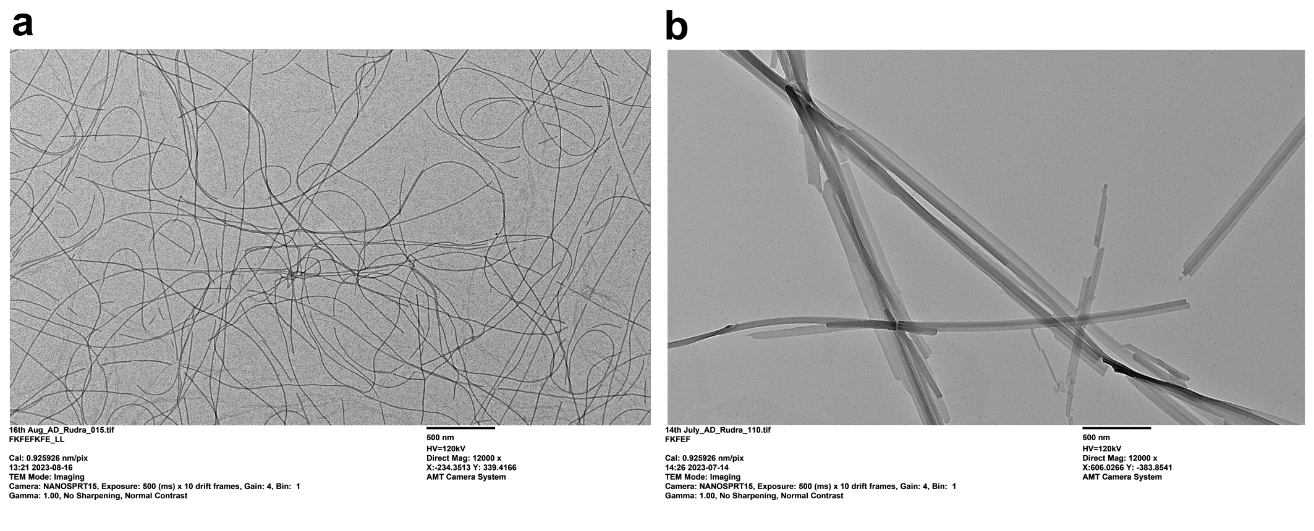
**Figure S9.** TEM images of (a) KFE8 and (b) KFE5 fibers.

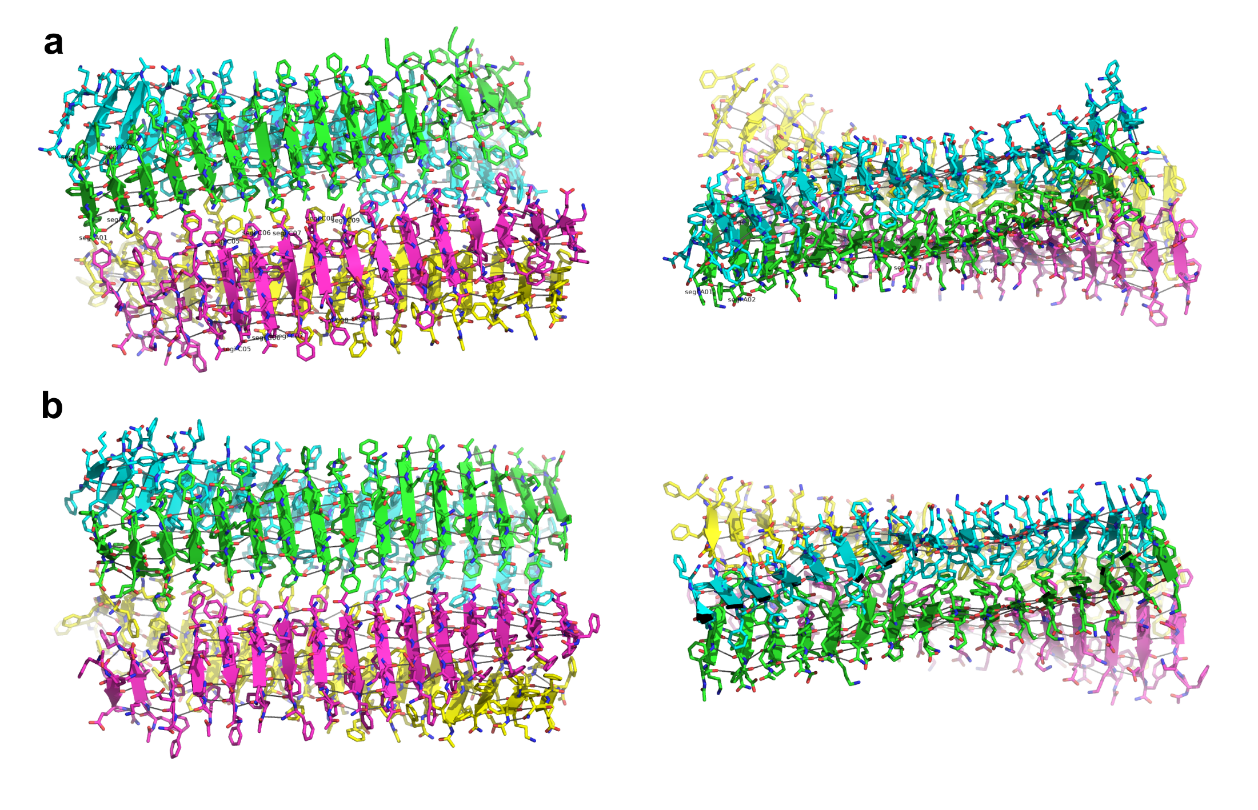
**Figure S10.** A 500 ns MD Simulation of the AlphaFold3 KFE5 model. The model is also based on the H. Wang KFE8 model. It consists of a sandwich of two anti-parallel β-sheet peptide assemblies, each comprising 16 strands that form a fiber. The model features an additional fiber that interacts via the hydrophobic side chains of the fiber edge (Phe:Phe). (a) At frame 14018. (b) At frame 24252.

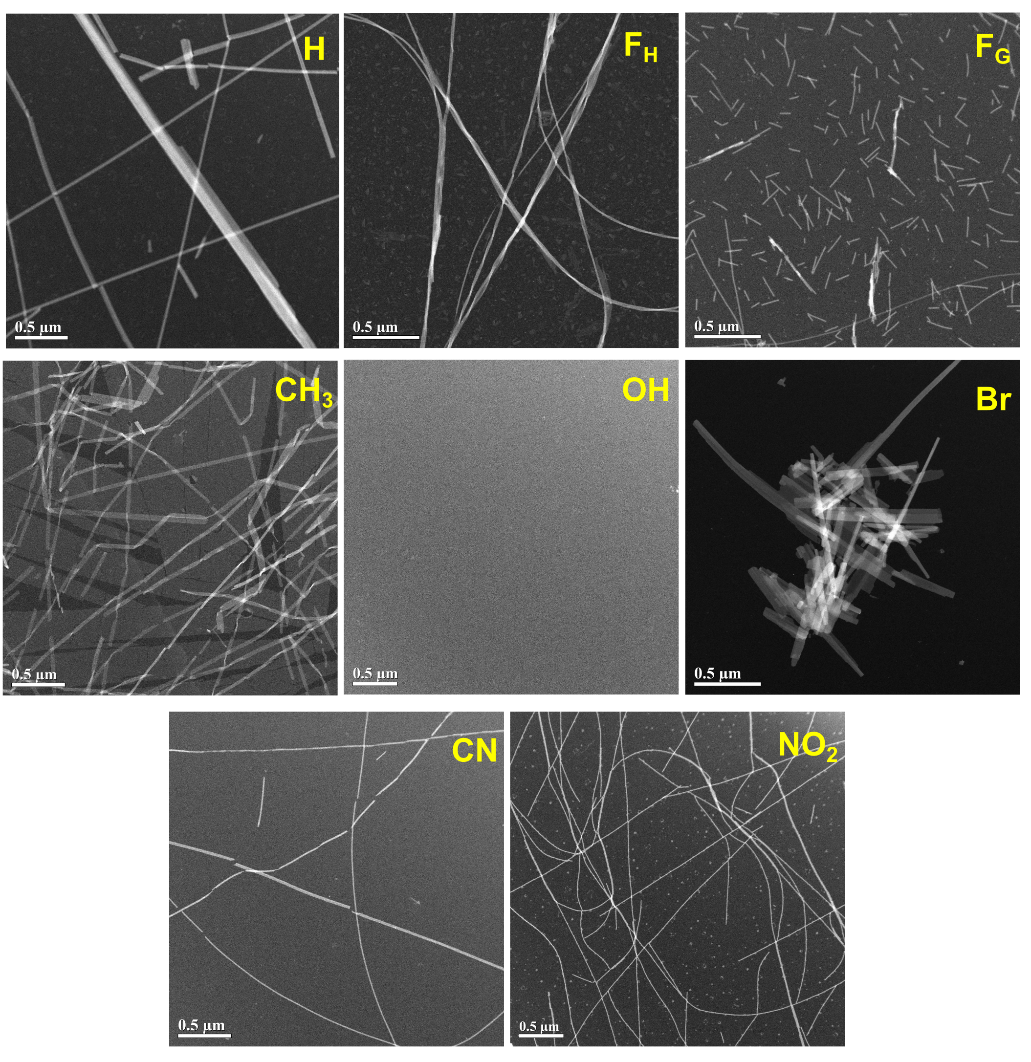
**Figure S11.** TEM images of KFE5 variant peptides. Scale bar is 0.5 µm.

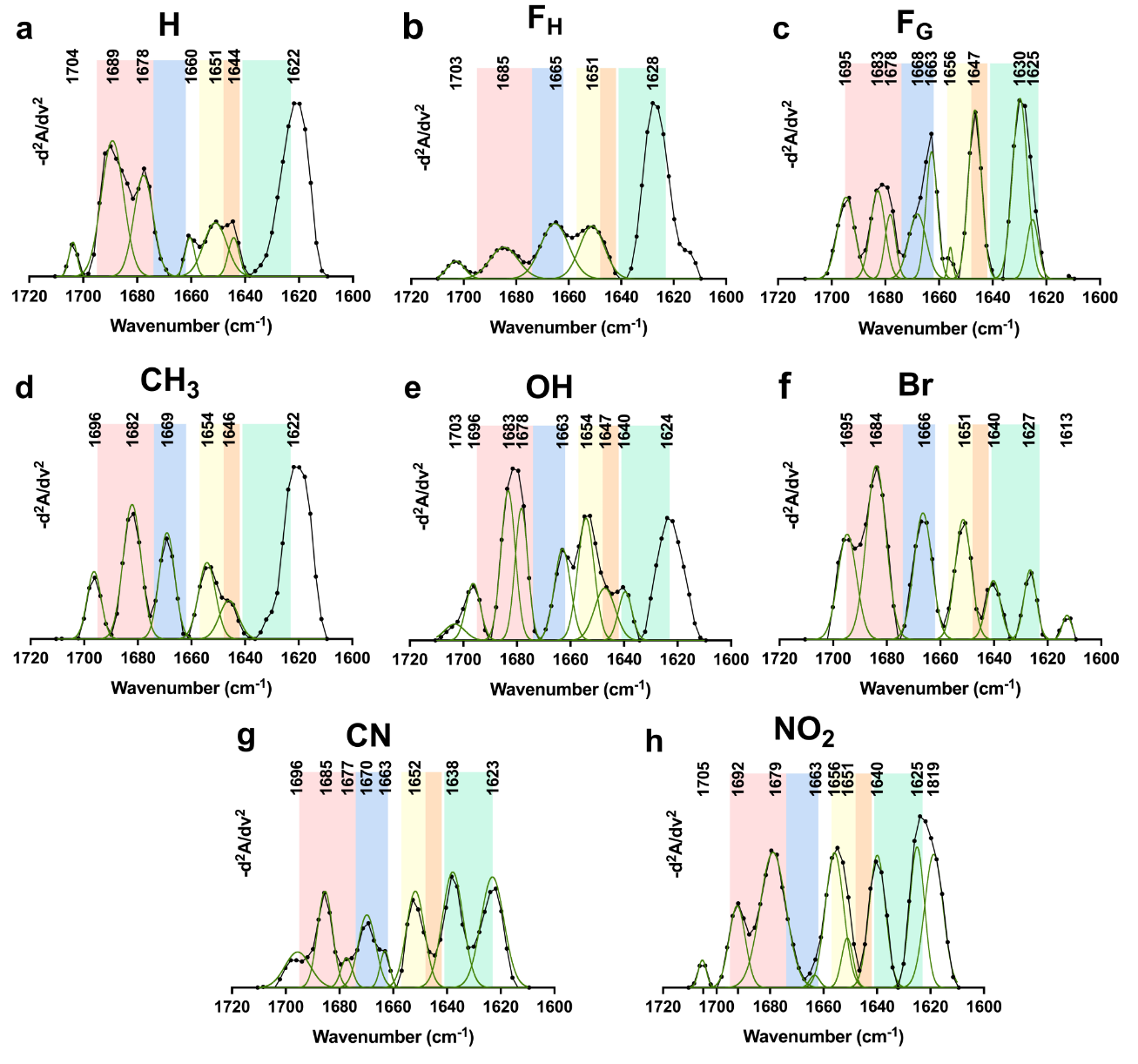

**Figure S12.** Second-derivative FT-IR spectra of KFE5 variants (a)-(h) in ultra-pure biological grade water.

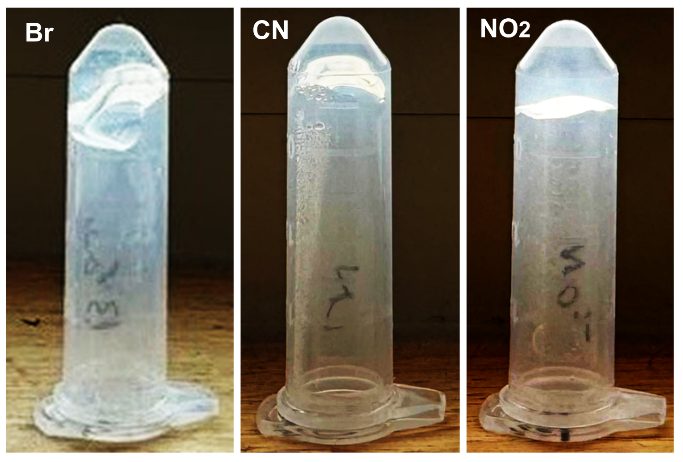
**Figure S13.** Representative images of gel formation by KFE5(Br) (2.5 mM), KFE5(CN) (5 mM), and KFE5(NO_2_) (5 mM) peptides.

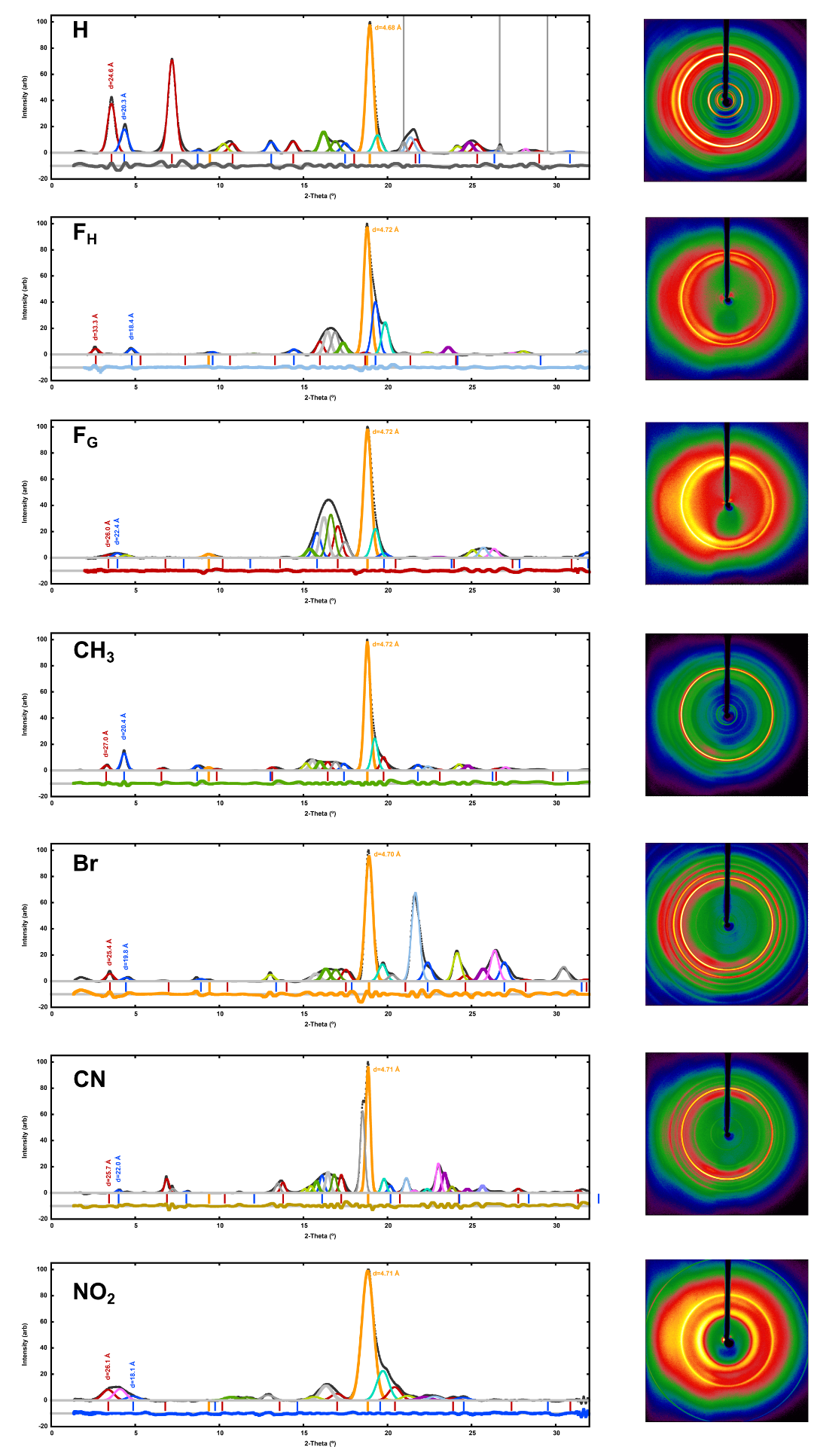

**Figure S14.** Fitting of the KFE5 XRD (powder diffraction) data along with its image (scan width 36°). Major peaks are (▬) β-sheet (d=4.693(3) Å), and (▬) helical repeat (d= 24.751(5) Å) with 4 higher-order peaks at d= 12.4, 8.25, 6.19, and 4.95 Å. Another (▬) major peak at d=20.38(2) Å has a possible (▬) (n=2) repeat at d=10.19 Å, (n=3) at d=6.79 Å, and n=4 at d=5.09 Å. Additional peaks occur at (▬) d=8.64(1), (▬) d=5.480(4), (▬) d=5.259(8), (▬) d=3.698(3), (▬) d=3.580(3), (▬) d=3.258(7). The (-) narrow peaks appear to be SiO_2_ from the capillary mounting glue. The (▬) residual is shown below. The scattering background was removed using an 18-parameter polynomial background fit in d1Dplot. The KFE5 sample was dissolved in methanol, pipetted into a MiTeGen capillary, and then left to dry prior to data collection. (right)

(a) The KFE5 diffraction image, from a 360° φ-scan.

(b) The KFE5(F_H_) XRD powder diffraction peak fitting, along with the diffraction image in the right panel.

(c) The KFE5(F_G_) XRD powder diffraction peak fitting, along with the diffraction image in the right panel.

(d) The KFE5(CH_3_) XRD powder diffraction peak fitting. along with the diffraction image in the right panel.

(e) The KFE5(Br) sample exhibits major peaks in common with KFE5. The β-sheet peak is at 2θ~19° (d=4.69 Å), the first helical twist peaks (d=24.7 Å) is seen at 2θ~ 3.5°, 7.13°, (10.7°), (14.4°), and 21.5°. The second helical twist (d=20.4 Å) is observed at 2θ~ 4.36° (8.7°), 13.0°, 17.5°. The large peak from 14.0° to 18.0° is due to the capillary holder and is much larger in the KFE5(Br) data due to smaller fraction of sample powder. The (▬) residual is shown below. The KFE5(Br) sample was dissolved in water, pipetted into a MiTeGen capillary, and then left to dry prior to data collection. (right) The KFE5(Br) diffraction image, from a 360° φ-scan.

(f) The KFE5(CN) XRD powder diffraction peak fitting. The main (▬) d=4.75(1) Å β-sheet peak is dominant. The filament structure has changed and is missing the strong (▬) d=24 Å feature. Two new peaks at (▬) d~13.41(1) Å and (▬) d~6.619(2) Å have appeared. Only two (▬) d~25.46(1) higher order peaks at n=2, d~12.7 Å and n=4, d~6.36 are possibly observed. The other helical (▬) d~19.36(3) Å reflections are possibly observed at n=4, d~ 4.84 Å, and n=5, d~ 3.87 Å. (right). The KFE5(CN) diffraction image, from a 360° φ-scan.

(g) The KFE5(NO_2_) XRD powder diffraction peak fitting, along with the diffraction image on the right panel.

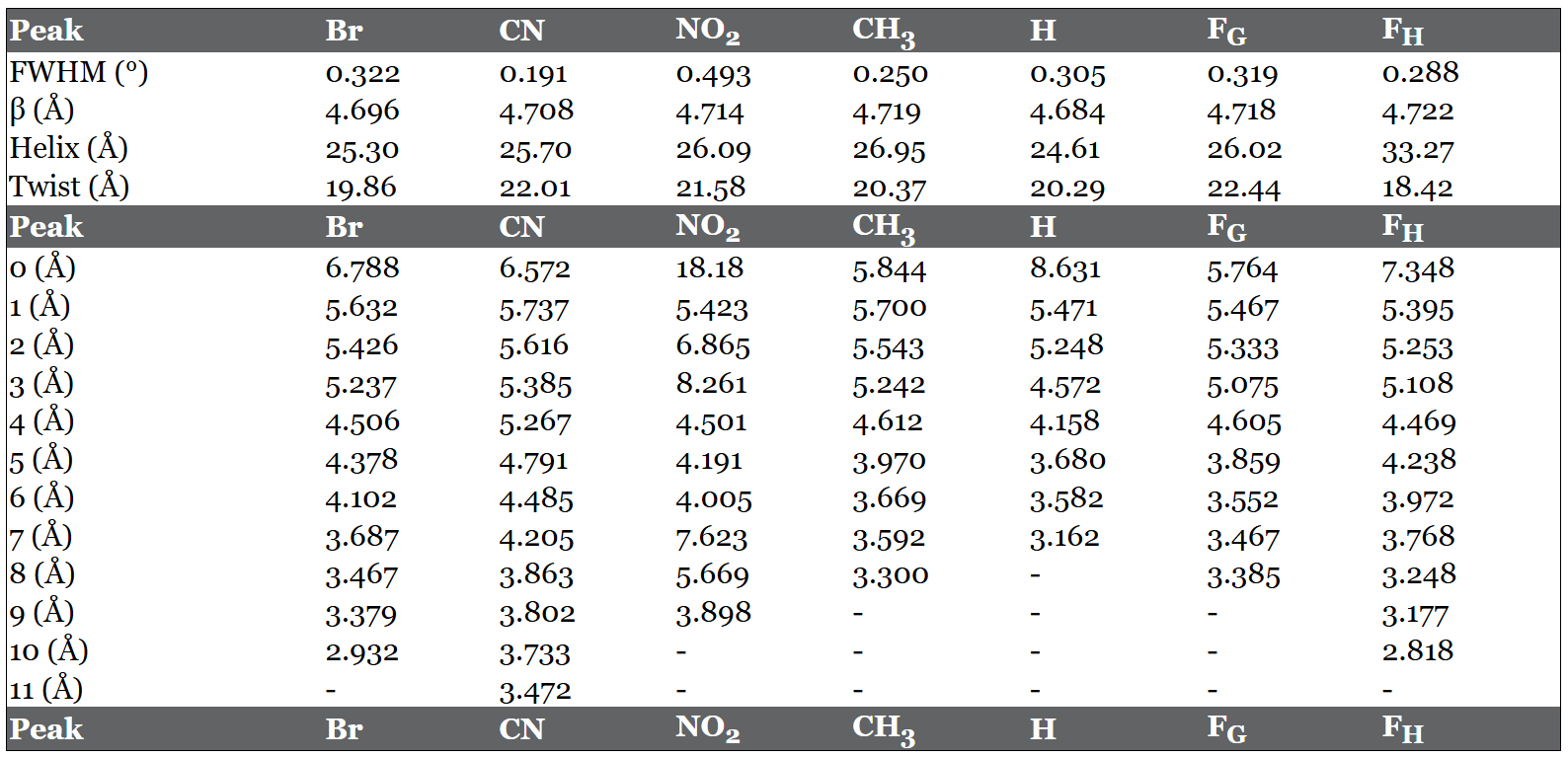
**Table S1:** WAXS peaks for 5-mer peptides

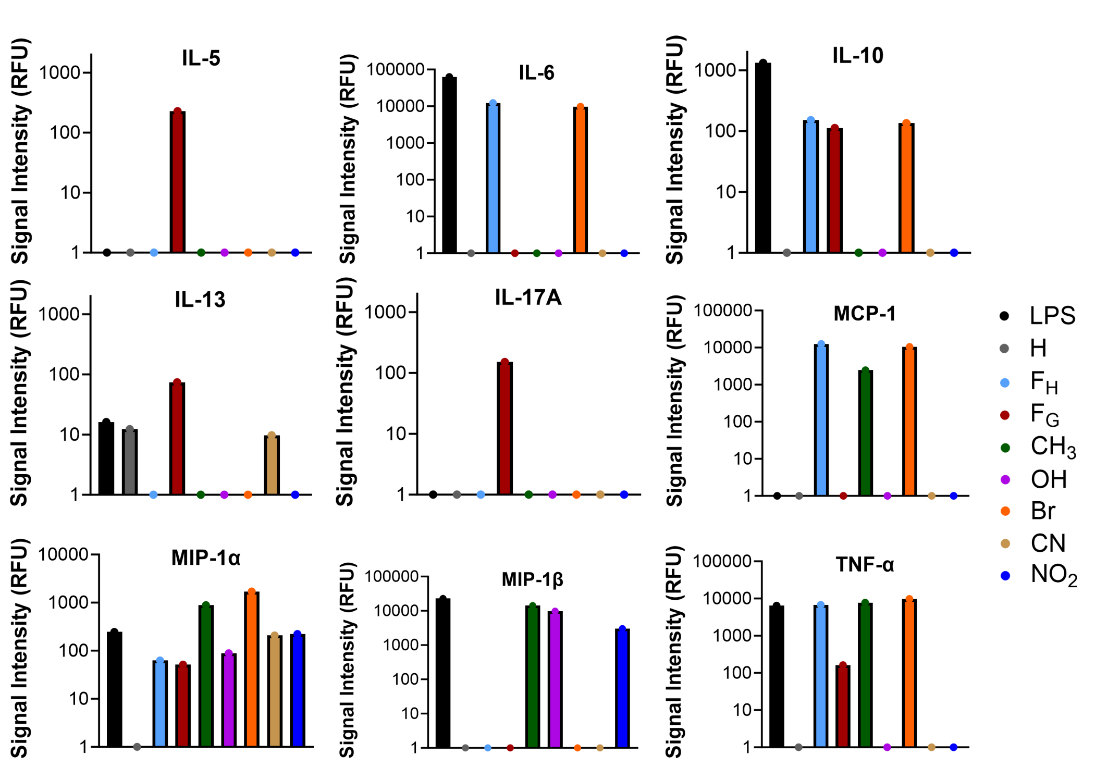
**Figure S15.** Cytokines and chemokines production in DCs treated with peptide nanofibers (10 μM) for 24h.

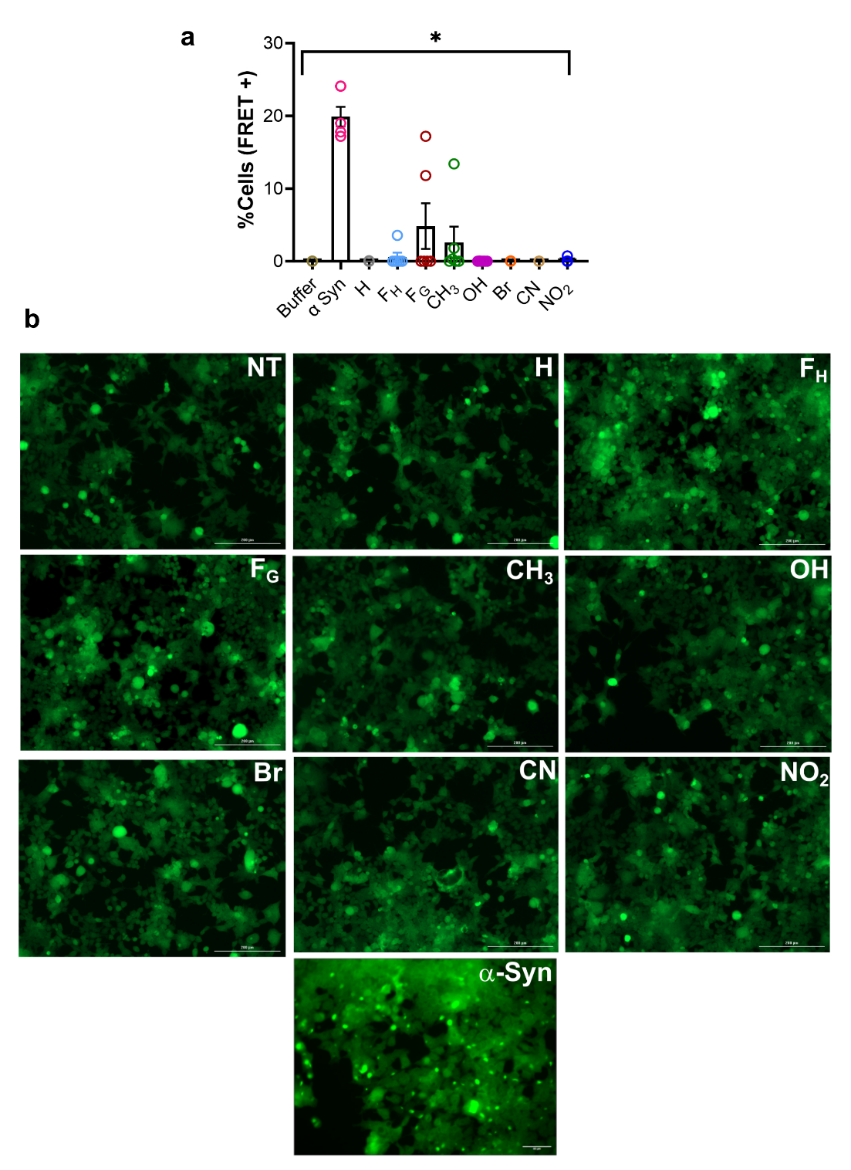
**Figure S16.** (a) % FRET positive cells as determined by the assay at 0.5 μM nanofiber concentration compared to α-synuclein PFFs (50 nM). (b) Microscopy images of 50 nM α-Syn and 0.5 μM 5-mer PNFs added to biosensor cells. Scale bars 200 µm. *p < 0.05, **p < 0.01, ***p < 0.001, ****p < 0.0001 as determined by a one-way ANOVA.

**Table S2:** Sequences and abbreviations for OVA conjugated peptides used in this study.

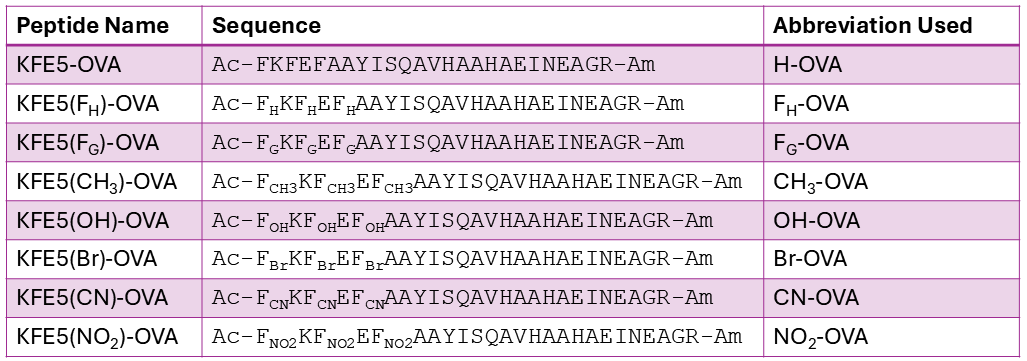

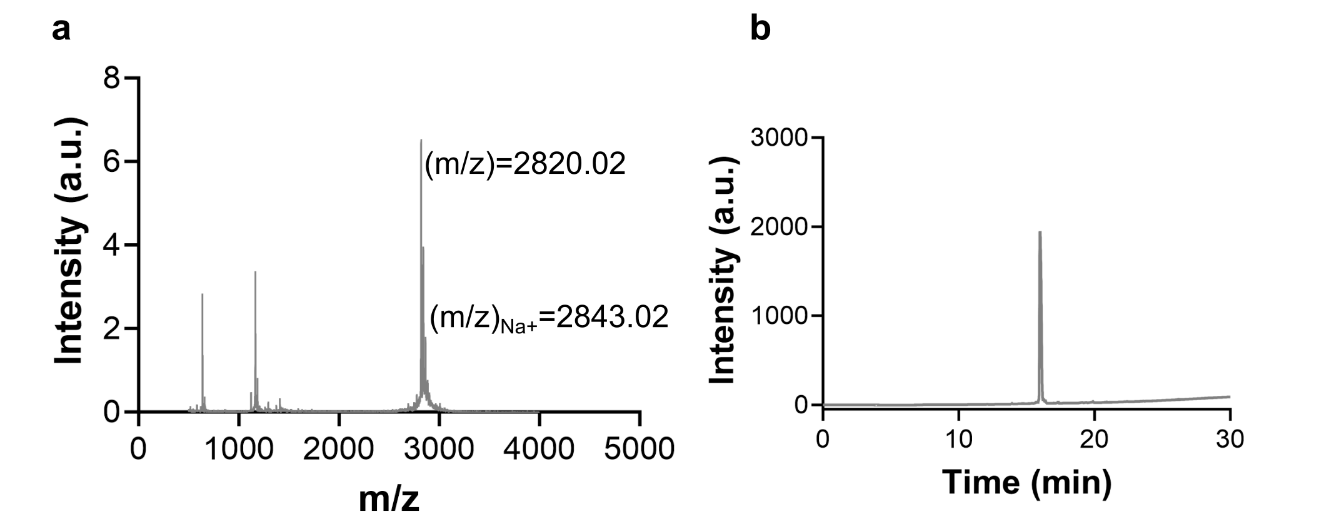

**Figure S17.** (a) MALDI-TOF and (b) HPLC profiles of KFE5-OVA

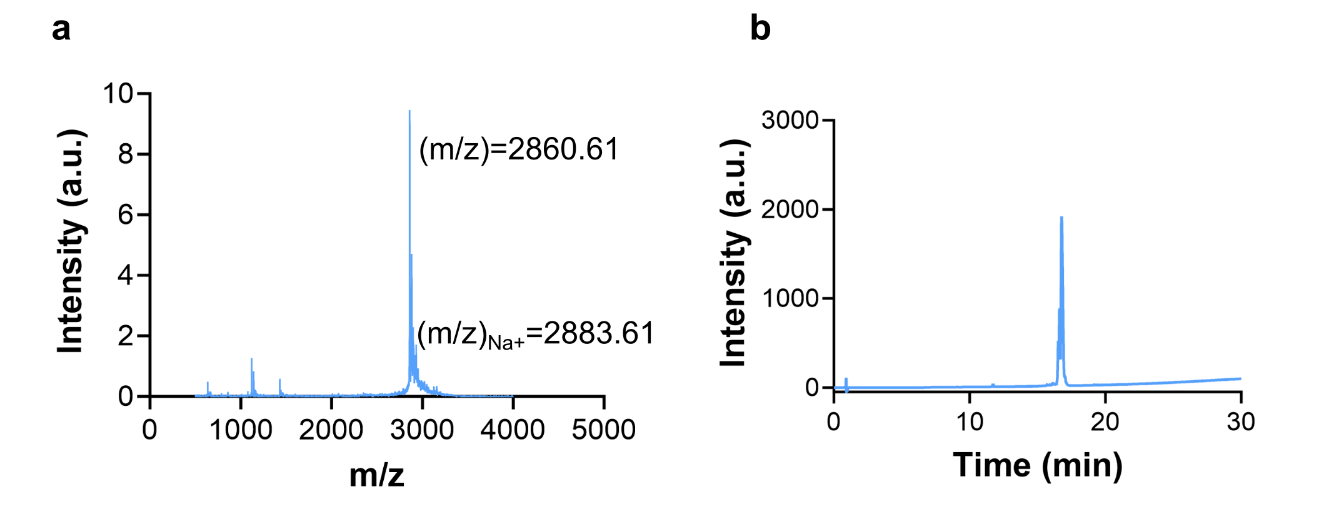

**Figure S18.** (a) MALDI-TOF and (b) HPLC profiles of KFE5(F_H_)-OVA

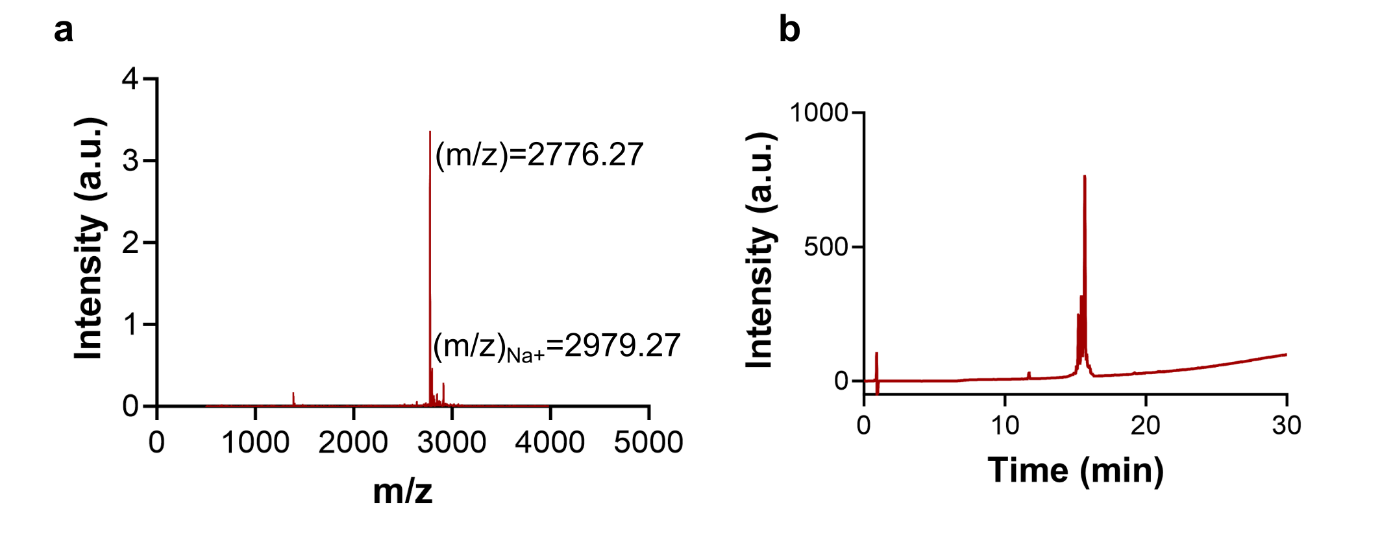

**Figure S19.** (a) MALDI-TOF and (b) HPLC profiles of KFE5(F_G_)-OVA

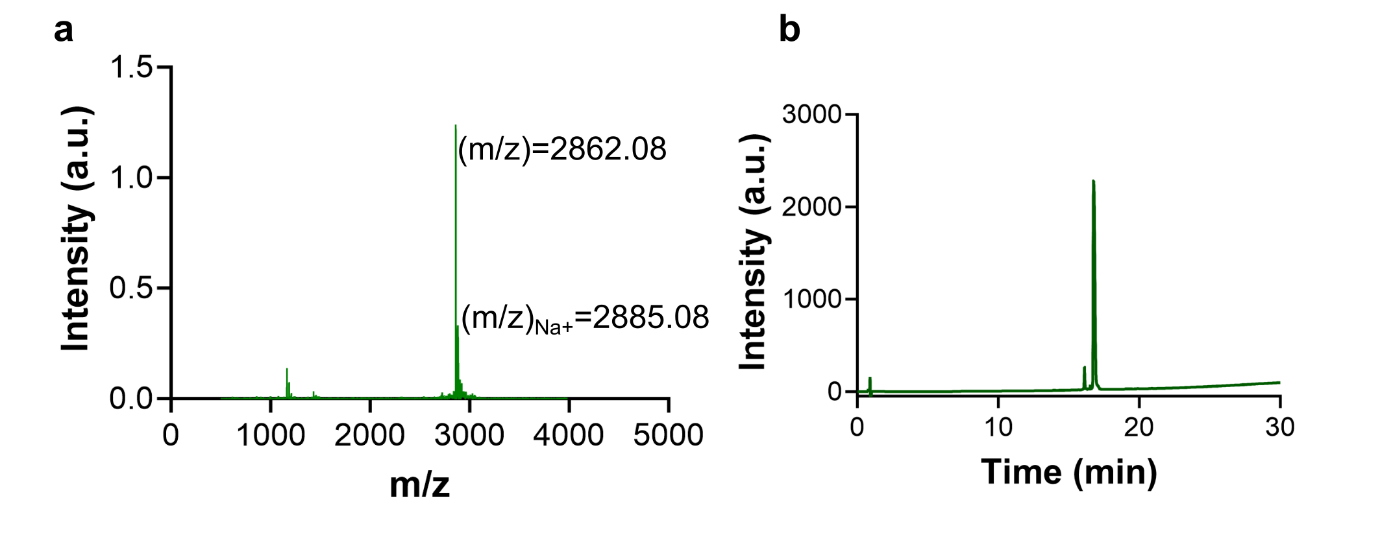

**Figure S20.** (a) MALDI-TOF and (b) HPLC profiles of KFE5(CH_3_)-OVA

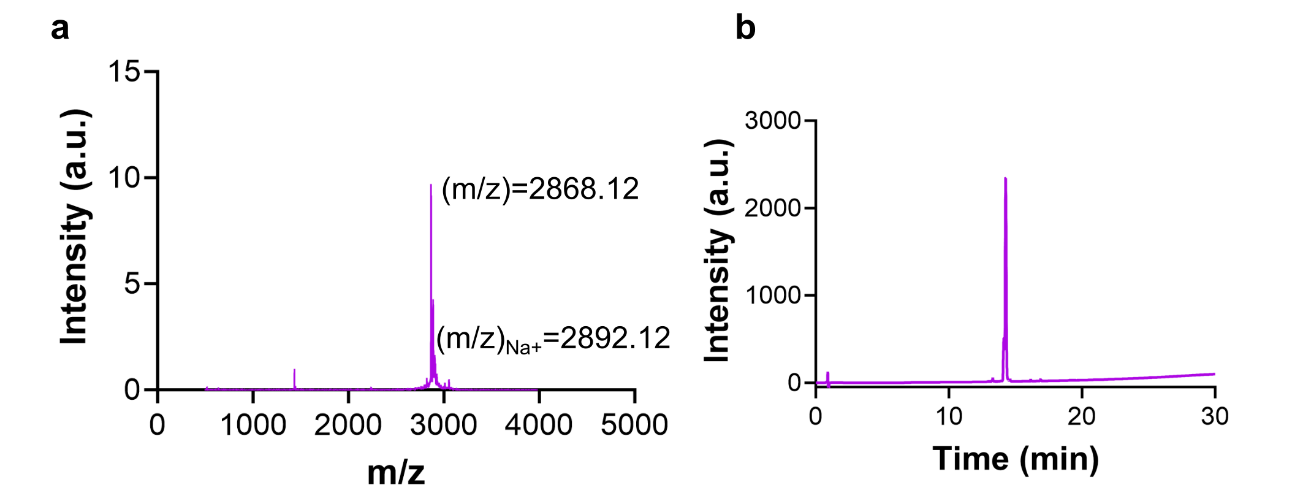

**Figure S21.** (a) MALDI-TOF and (b) HPLC profiles of KFE5(OH)-OVA

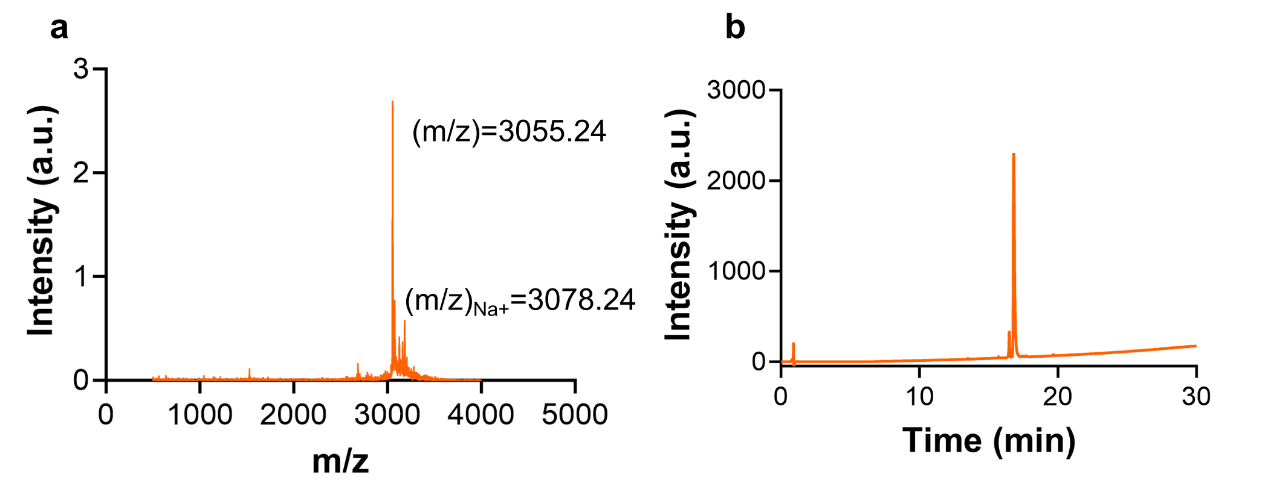

**Figure S22.** (a) MALDI-TOF and (b) HPLC profiles of KFE(Br)-OVA

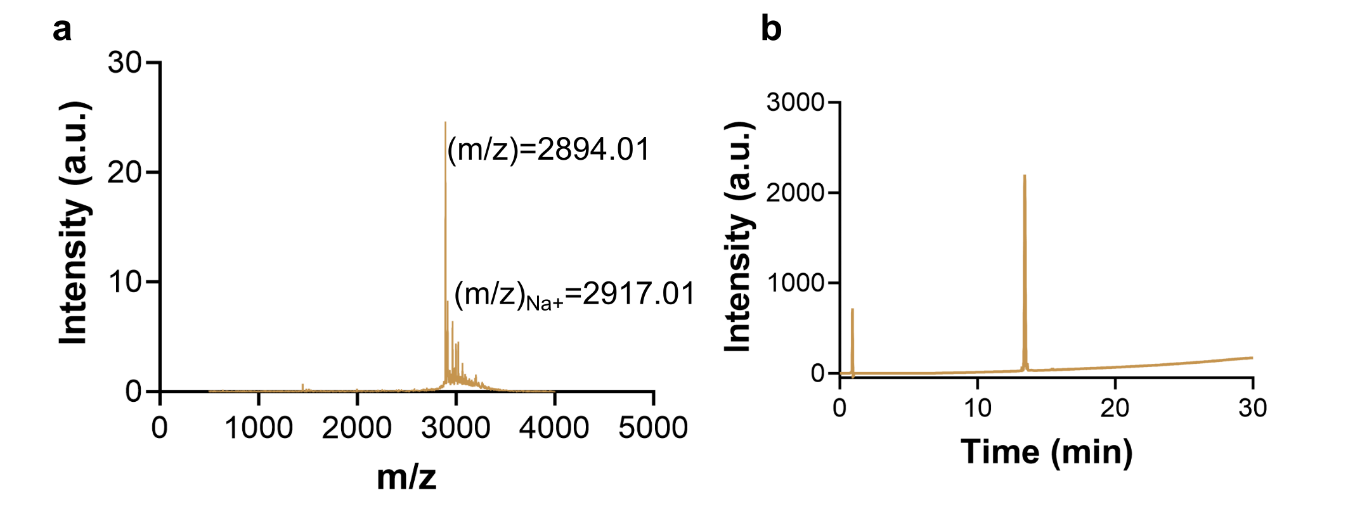

**Figure S23.** (a) MALDI-TOF and (b) HPLC profiles of KFE5(CN)-OVA

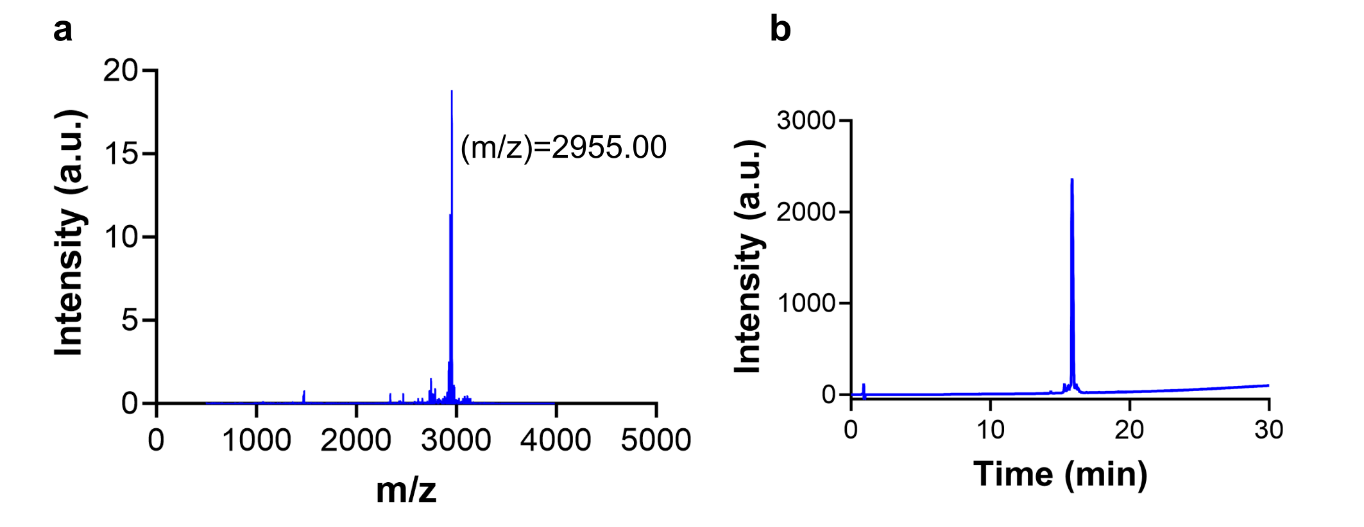

**Figure S24.** (a) MALDI-TOF and (b) HPLC profiles of KFE5(NO_2_)-OVA

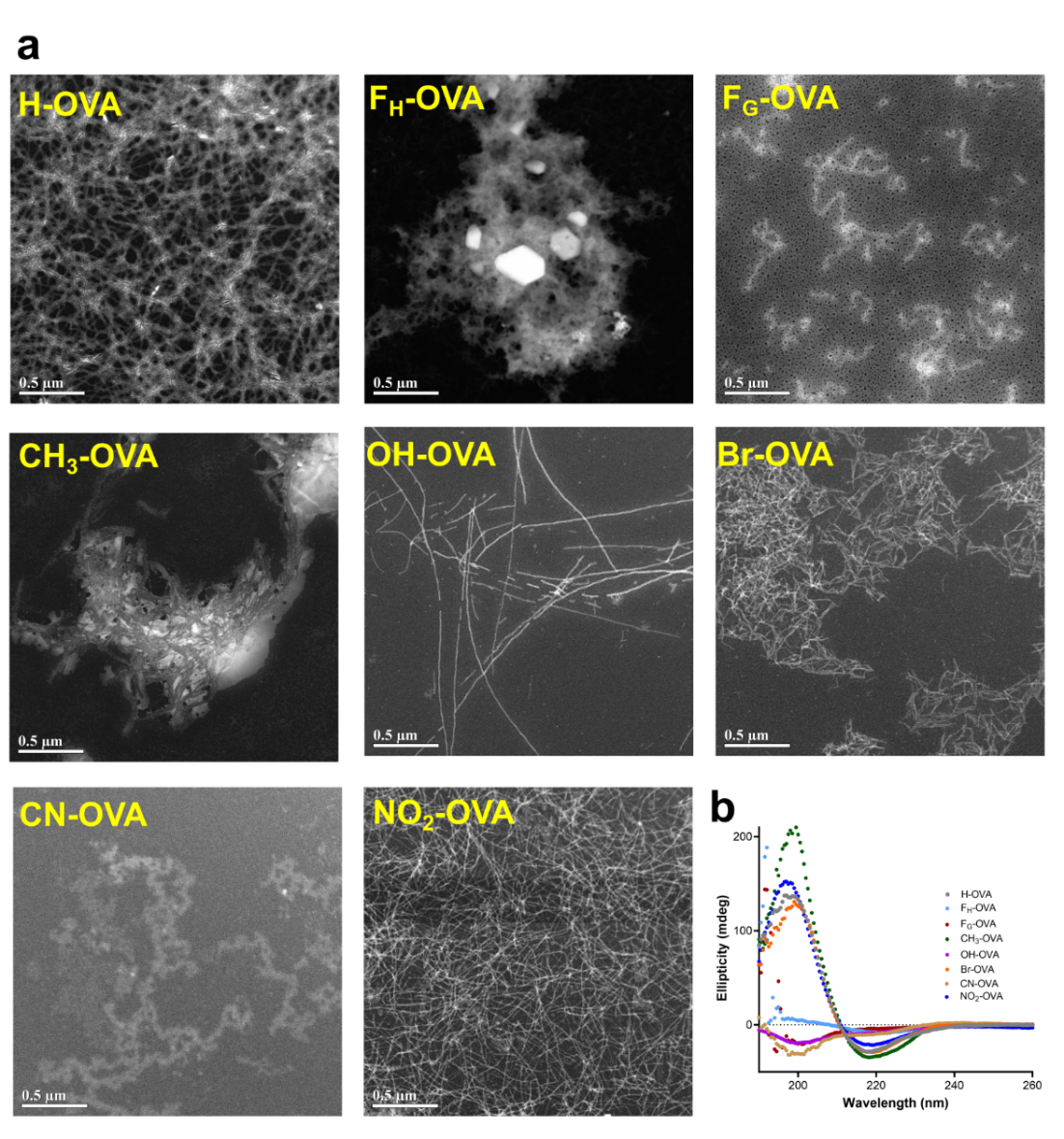
**Figure S25.** (a) TEM images of self-assembled OVA-conjugated pentapeptides as marked in the figure. (b) Respective CD profiles of the fibrils.

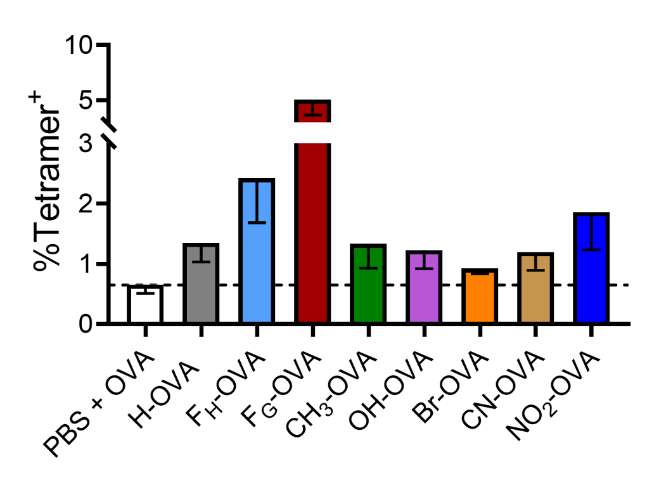

**Figure S26.** The frequency of OVA-specific CD4^+^ T cells is represented as a percentage of total CD4^+^ T cells in the spleens of vaccinated mice.
